# Apusomonad rhodopsins, a new family of ultraviolet to blue light absorbing rhodopsin channels

**DOI:** 10.1101/2025.04.02.646541

**Authors:** Luis Javier Galindo, Shunki Takaramoto, Takashi Nagata, Andrey Rozenberg, Hiroto Takahashi, Oded Béjà, Keiichi Inoue

## Abstract

Apusomonads are a clade of understudied sediment-dwelling bacterivorous protists sister to Opisthokonta. Recently, apusomonads have been found to show a negative phototactic response to blue light. Here, by screening available apusomonad ‘omics data we found genes of a distinct group of microbial rhodopsins, the ApuRs. ApuRs heterologously expressed in mammalian cells absorbed near-UV/violet or blue light, suggesting that ApuRs could be involved in apusomonads’ photoavoidance response. Electrophysiological measurements indicate that ApuRs are anion-selective rhodopsin channels which evolved independently of the family of channelrhodopsins widespread in other unicellular eukaryotes. Among the known rhodopsin channels, ApuRs collectively demonstrate the most blue-shifted absorption spectra. In ApuRs, the channel opening is triggered by photoisomerization of the retinal from its all-*trans* form to 13-*cis* and 11-*cis* forms. We found that intracellular proton transfer is involved in channel opening and determined the channel’s open/close kinetics. These findings expand our understanding of the photobiology of heterotrophic flagellates and showcase the significance of these organisms as a source of new rhodopsin families with unanticipated functions.

## Introduction

Light is an essential source of energy for life and one of the most important sources of information about the external world ^1^. Thus, organisms have evolved to perceive and respond to it, leading to its role as a regulator of a wide range of cellular processes (e.g., phototaxis, growth, and metabolism) ^1–7^. These light-driven cellular processes are ultimately linked to the organism’s capability to survive and adapt to its environment. The main players in cellular light perception systems are photoreceptor proteins, which have evolved to respond to the spatial, temporal and spectral variations of light ^1,7,8^. A particularly striking diversity is demonstrated by photoreceptors among microbial rhodopsins ^9^. While the most abundant rhodopsins are involved in harvesting sunlight as a source of energy by prokaryotic photoheterotrophs ^10^, many microbial rhodopsin families function as light sensors utilizing a range of mechanisms: from transducer activation in prokaryotic rhodopsin sensors ^11,12^ to changing membrane potential in channelrhodopsins ^13,14^ and directly regulating cellular signaling in enzymerhodopsins ^15–17^.

Unicellular eukaryotes (protists) have been shown to harbor diverse families of microbial rhodopsins ^13,18–21^. Some of protist microbial rhodopsins have also generated wide interest due to their biotechnological applications, particularly in the field of optogenetics ^20,22–26^. The first and best known example is channelrhodopsin-2 (*Cr*ChR2) from the unicellular green alga *Chlamydomonas reinhardtii*, which has been used to depolarize and activate mammalian neurons ^13,24,27,28^. Since the discovery of channelrhodopsins in *Chlamydomonas*, many more microbial rhodopsins from protists have been found, and some have been successfully adapted as optogenetic tools^15,20,25,29^.

Three rhodopsin groups coming exclusively from unicellular eukaryotes have attracted particular attention in recent decades: channelrhodopsins, enzymerhodopsins and eukaryotic rhodopsin pumps. Despite progress made in the biophysical characterization of these families, thanks to the optogenetic potential of some of these proteins ^30–32^, experimental evidence of their physiological roles is available only from studies on very few unicellular eukaryotes. Examples include channelrhodopsins involved in phototaxis in *C. reinhardtii* ^13,14^, rhodopsin proton pumps with divergent roles in photosynthesis in marine diatoms ^30,33^, proton pumps involved in acidification of food vacuoles in the heterotrophic dinoflagellate *Oxyrrhis marina* ^34^ and a rhodopsin-guanylyl cyclase fusion in the zoosporic fungus *Blastocladiella emersonii* responsible for phototactic behavior in its zoospores ^15^. What is more, these studies become more scarce as we look towards the highly diverse group of heterotrophic flagellates.

Heterotrophic flagellates (HFs) are colorless, non-photosynthetic small predatory protists feeding mainly on bacteria or other protists and are likely to be the most abundant eukaryotes on Earth ^35^. Moreover, every major eukaryotic group seems to have evolved from a HF lineage ^36^. In the different evolutionary reconstructions of the origins of eukaryotes, a lineage of heterotrophic flagellates is always found near the root of eukaryotes ^37–39^, suggesting that LECA (Last Eukaryotic Common Ancestor) resembled extant HFs ^40^. Despite their ecological and evolutionary importance, HFs remain highly understudied and new species are constantly being discovered ^41–43^. Accordingly, our knowledge about light sensing and rhodopsin diversity in HFs is limited and experimental efforts in this regard have been concentrated on biophysical characterization of the diverse repertoire of channelrhodopsins found in stramenopiles (cation channels, including potassium channelrhodopsins or KCRs, in placidids, bicosoecids and a hyphochytriomycete ^44–46^ and anion channels in labyrinthulids ^47^, bicosoecids ^48^, MAST-3 ^49^ and MAST-4 ^50^) and occasionally in other HFs (*Goniomonas* ^49^, *Colponema* ^44^, putative uncultured katablepharids ^44,51^), rhodopsin-phosphodiesterase fusions (Rh-PDEs) in choanoflagellates ^17^ and the non-sensory proton pumps in the non-photosynthetic dinoflagellate *O. marina* ^52^. Despite the modest sample of the characterized proteins, some of these rhodopsins, such as MerMAIDs ^50^, RubyACRs ^47^ and Rh-PDEs ^53^ were found to have an optogenetic potential, and KCRs ^45,46^ are becoming a tool of choice for neuron inhibition. Characterization of new microbial rhodopsins from HFs presents a valuable opportunity to understand the ecological dynamics and evolution of eukaryotic life, as well as to search for new biotechnologically useful rhodopsins.

The apusomonads (Apusomonadida) are bacterivorous biflagellate HF protists that are found gliding on freshwater and marine sediments all over the world ^54–57^. They are the sister lineage to Opisthokonta, which includes animals, fungi and their unicellular relatives. Thus, apusomonads retain ancestral features crucial to understanding early opisthokont evolution ^57–60^. Recently, two large efforts managed to isolate in culture a representative sample of diverse apusomonads, including four new genera and seven new species ^58,61^. To resolve phylogenetic relationships among apusomonads and to study key traits through comparative genomics, these studies generated transcriptomic data for the new species ^57^.

Here we aimed to unveil some of the hidden diversity of microbial rhodopsins from HFs by screening all available apusomonad omics data, including the new transcriptomic and environmental metagenomic data. Our screening efforts led to the discovery of twelve full-length microbial rhodopsins belonging to an entirely new family unique to apusomonads.

## Results

### A new class-level group of microbial rhodopsin from apusomonads

By analyzing all of the available genome and transcriptome assemblies of apusomonads ^57^, we discovered that half of them coded for proteins from a distinct group of microbial rhodopsins which we named apusomonad rhodopsins (ApuRs). A search for closely related proteins in assemblies from other unicellular creatures yielded no results indicating that ApuRs are specific to apusomonads. While apusomonads are known as benthic protists, ApuRs were surprisingly found in the pelagic environmental data from the *Tara* Oceans metagenomes and metatranscriptomes ^62,63^: one ApuR type closely related to *Tt*ApuR from *T. trahens* and a distinct type found predominantly at station TARA_39 in the North Indian Ocean (represented in our dataset by *Tara*ApuR derived from the MERC assembly ^64^). By combining all of the different sources, we collected a total of twelve non-redundant ApuR proteins with intact rhodopsin domains and a Lys in transmembrane helix TM7 (Table 1). The highest number of ApuRs genes was found in *Singekia* sp. FB-2015 which in addition to three intact genes, also has a putatively photoinactive ApuR with a His residue in TM7 in place of the Lys. The proteins belonging to the family demonstrate a high sequence diversity with pairwise sequence identities in the region covering the transmembrane part of the protein ranging from 60% between the sister sequences *Sf*ApuR1 and *Sf*ApuR2 from *Singekia franciliensis* to as low as 24% between *Pk*ApuR from *Podomonas kaiyoae*, representing the most distinct sequence, and other ApuRs. The amino acid sequence motifs in TM3, usually conserved in functionally related ion-transporting microbial rhodopsins, were also found to diverge, although their diversity could be fit into two classes: most of the sequences demonstrate the DXQ motif (where X is Ala or Ser) and a minority – the XTQ motif (where X is Ala, Met, Ile or Cys) (see Table 1).

**Table 1.** General information about ApuRs investigated in this study.

| **Apusomonad species** | **ApuR** | **TM3 motif** | **HsBR D212** | **$\lambda_{\max}$ (nm)** |
| --- | --- | --- | --- | --- |
| *Chelonemonas dolani* | CdApuR<sub>DAQ</sub> | DAQ | D | 385 |
| *Thecamonas trahens* | TtApuR<sub>DAQ</sub> | DAQ | D | 385 |
| *Karpovia croatica* | KcApuR<sub>DAQ</sub> | DAQ | D | 369 |
| *Singekia franciliensis* | SfApuR1<sub>DSQ</sub> | DSQ | D | 452 |
|  | SfApuR2<sub>DAQ</sub> | DAQ | D | 386 |
| *Singekia montserratensis* | SmApuR<sub>DAQ</sub> | DAQ | D | 391 |
| *Singekia* sp. FB-2015 (AF-17) | SsApuR1<sub>ATQ</sub> | ATQ | E | 489 |
|  | SsApuR2<sub>DAQ</sub> | DAQ | D | 386 |
|  | SsApuR3<sub>ITQ</sub> | ITQ | S | N.D. |
| *Mylnikova oxoniensis* | MoApuR<sub>MTQ</sub> | MTQ | E | 449–485 |
| *Podomonas kaiyoe* | PkApuR<sub>CTQ</sub> | CTQ | T | 398 |
| **Environmental transcriptome** | **ApuR** | **TM3 motif** | **HsBR D212** | **$\lambda_{\max}$ (nm)** |
| ERR1711916<br>(Tara Oceans environmental dataset) | TaraApuR<sub>DAQ</sub> | DAQ | D | N.D. |

### Phylogeny of microbial rhodopsins

Our phylogenetic tree of the diversity of microbial rhodopsins recovered the monophyly of ApuRs with a high support value with the ApuR clade branching off deeply among other microbial rhodopsins (Fig. 1). Despite lack of strong support for the deepest branches, ApuRs do not appear to have affinity with any other rhodopsin family (see Fig. 1). Nearly half of the apusomonad species for which sufficient genetic data were available were found to possess ApuR genes (Table 1 and Fig. 1b). Gene tree-species tree reconciliation, under the most parsimonious scenario, places the root of the ApuR family in the *Podomonas*–*Mylnikova* clade with a subsequent acquisition by an ancestor of the Thecamonadinae (Fig. S1). Furthermore, the gene phylogeny strongly suggests that the TM3 motif ancestral for the ApuR family was XTQ which evolved to DAQ in one of the branches of the gene tree with the DAQ ApuRs appearing solely among the Thecamonadinae species (Fig. 1b). To explain the appearance of the earlier diverging XTQ genes in the thecamonadine *Singekia* sp. FB-2015, alongside *Ss*ApuR2_DAQ_, an ancient gene duplication is hypothesized with several subsequent gene loss events (see Fig. S1). An alternative explanation for this apparent incongruence is a transfer of an XTQ gene to the lineage of *Singekia* sp. FB-2015 from an unsampled or extinct apusomonad lineage.

**Figure 1.**
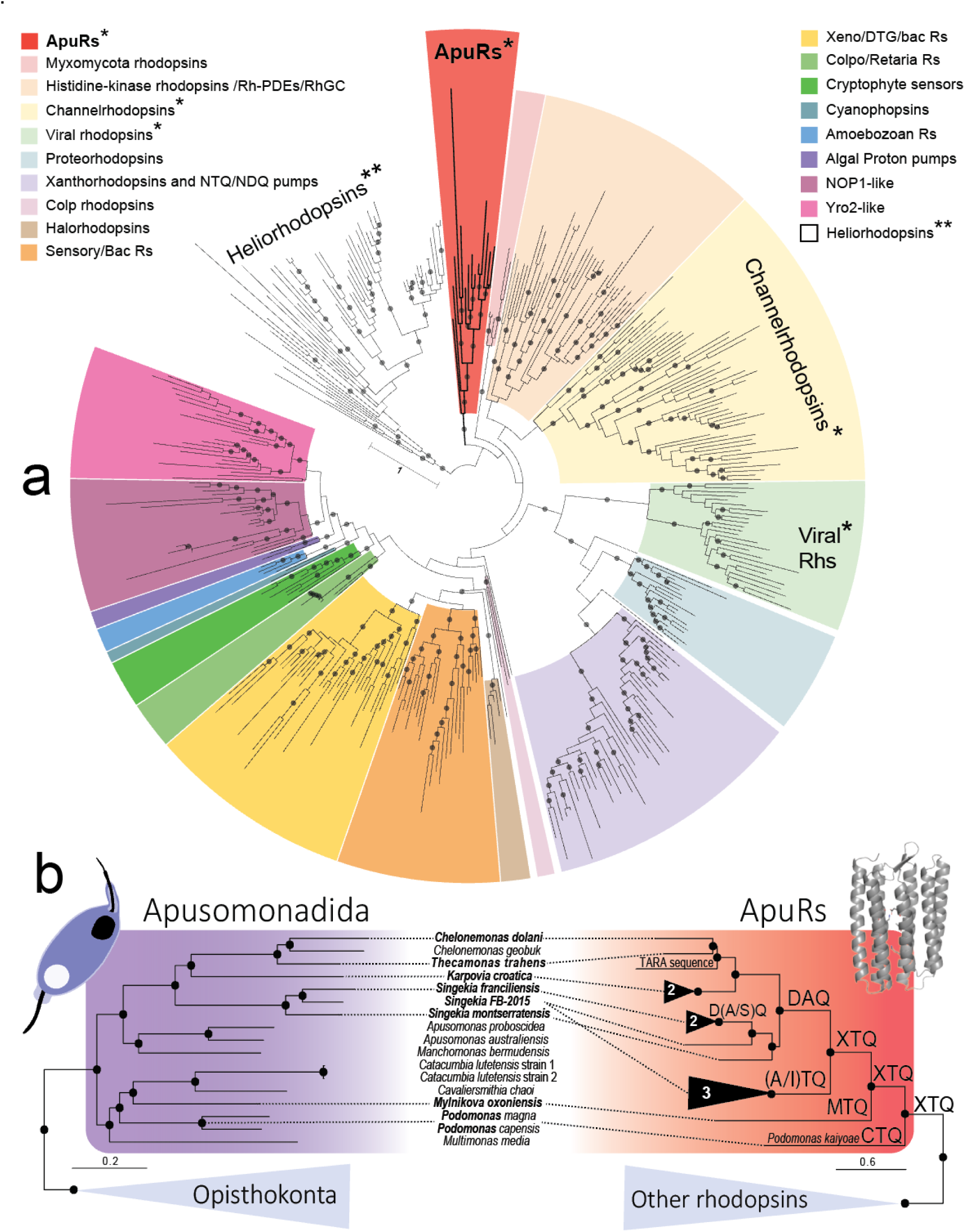
Phylogeny and evolution of microbial rhodopsins and ApuRs. **a** Maximum likelihood (ML) phylogenetic tree of the diversity of microbial rhodopsins. Microbial rhodopsin families are highlighted in different colors. The phylogenetic dataset is a modified version of the dataset by Rozenberg et al. 2021^9^, enriched for HF-Rhs (including ApuRs) and now including 396 sequences. The phylogenetic tree was reconstructed with the Q.pfam+F+R8 model as chosen per BIC. Statistical support was generated with 1000 ultrafast bootstraps (ufbs). This tree is rooted in a selection of heliorhodopsins as outgroups (non-highlighted branches). **b** Graphical comparison of a ML phylogenomic tree of Apusomonadida (left; 303 conserved proteins; LG+C60+G/PMSF)^57^ vs a simplified version of the same microbial rhodopsin ML phylogeny highlighting the diversity of ApuRs (right). Black dots indicate statistical support values of ≥ 95% ufbs. A single asterisk (*) indicate microbial rhodopsin clades in which proteins mainly exhibit channeling activity and double asterisks (**) indicate clades in which some proteins exhibit channel activity.

The narrow distribution of ApuRs solely among Apusomonadida is reminiscent of the pattern of some families of channelrhodopsins, such as the strict appearance of the green algal cation channelrhodopsins (CCRs) and the related prasinophyte anion channelrhodopsins (ACRs) associated with the eyespot and vision in green algae ^65^. This might imply that ApuRs similarly underwent adaptation to a specific physiological role or cellular localization. Given the lack of morphologically discernible eyespots in apusomonads, physiological experiments would be required to correlate the presence of ApuRs with phototactic behavior.

### Structural features and conserved residues in ApuRs

The predicted structures of ApuRs demonstrate the architecture of regular microbial rhodopsins with seven transmembrane helices, the N-terminus facing the extracellular side, a π-bulge in transmembrane helix 7 (TM7) with a conservative Lys residue bound to the all-*trans*-retinal moiety, in addition to a C-terminal part of variable length (Fig. 2). The highest structural similarity to ApuRs was observed for rhodopsins from the bacteriorhodopsin-like families, *Leptosphaeria* rhodopsin (LR) and schizorhodopsin SzR4 (Fig. S2). Incidentally, the proteins in these families are either known to form trimers or are strongly suspected to do so. To explore whether ApuRs might be forming trimers as well, we predicted homoligomers with varying numbers of monomers for ApuRs and reference rhodopsins. Generally, for rhodopsins with known oligomeric structures the predictions corresponding to their natural oligomeric number received the highest interface predicted template modelling (ipTM) scores, although for BCCRs and some pentameric rhodopsins, dimeric and trimeric predictions, respectively, appeared as close second. For all of the predicted ApuR structures, only trimeric predictions yielded sufficiently high ipTM scores exceeding or approaching 0.8, except for *Ss*ApuR2 for which all the predictions were low-scoring (Fig. S3a). This strongly suggests that ApuRs form trimers. A closer look at the trimeric structures revealed that they match that of the bacteriorhodopsin from *Halobacterium salinarum* (*Hs*BR) (Fig. S3b). The relatively high structural similarity between ApuRs to the trimeric rhodopsins (except BCCRs), coupled with the lack of a clear phylogenetic affinity to them, indicate that this similarity is plesiomorphic: inherited from a common ancestor of regular microbial rhodopsins. On the other hand, TM-scores in comparisons to structures of known groups of rhodopsin channels, such as dimeric and trimeric channelrhodopsins, bestrhodopsin *Tara*-RRB and viral rhodopsins (VRs) were much lower (see Fig. S2).

**Figure 2.**
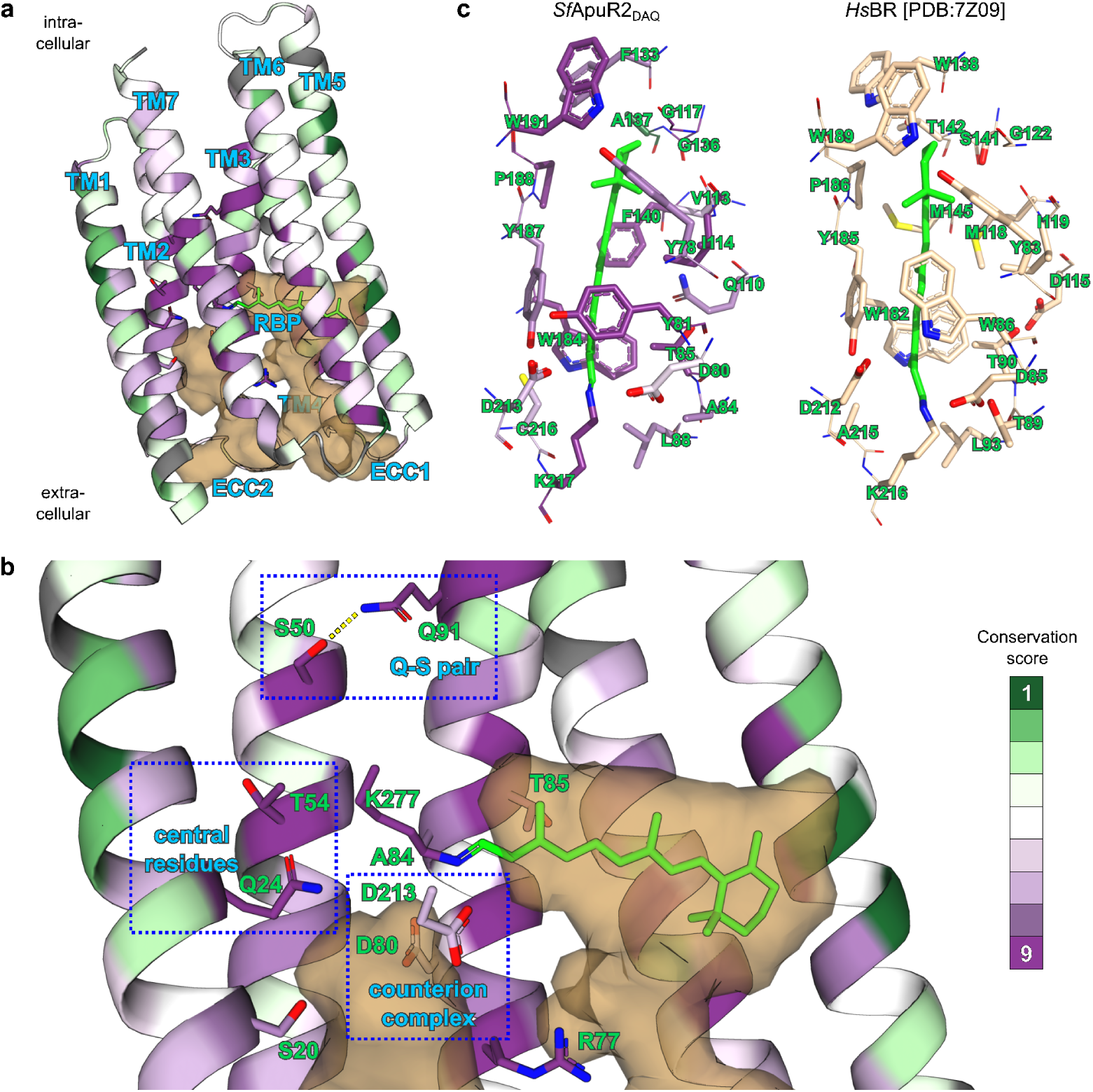
Structural features of ApuRs exemplified by the predicted structure of *Sf*ApuR2. **a** The overall structure of ApuRs with residues colored according to Consurf conservation score. Three interconnected cavities are indicated as red volumes: the retinal binding pocket (RBP), extracellular cavity 1 (ECC1) above it and extracellular cavity 2 (ECC2) discernible only in some ApuR structures. Transmembrane helices TM1-7 are indicated as well. Most of the TM6 and TM7 helices are removed for clarity. Extracellular side above. **b** A zoom-in on the region around the retinal Schiff base with conserved non-hydrophobic residues and residues of the TM3 motif indicated and labelled. **c** Detail of the residues forming the retinal binding pocket in *Sf*ApuR2 and their homologs in *Hs*BR. View from the extracellular side.

Only one of the ApuRs, *Ss*ApuR2, was predicted to have a signal peptide although there is uncertainty in the position of the start codon in the corresponding transcript. As is usually the case in other families of microbial rhodopsin, most of the structural variation between ApuRs is concentrated in the intra- and extracellular loops between the transmembrane helices (ICLs and ECLs) and the C-terminal region. While ICL3 (EF loop) shows little sequence conservation and is highly variable in length, ECL1 (BC loop) was found to accept one of two main configurations (Fig. S4a). The putatively ancestral structure of ECL1 contains a short antiparallel β-sheet similar to that of proteins from the bacteriorhodopsin-like superclade, while the putatively derived shorter versions of ECL1 in all ApuRs from *Singekia* spp. lack the β-sheet entirely. An analogous evolutionary pattern with alternative ECL1 types has been reported for the family of fungal rhodopsins ^66^. Similarly to ChRs, the C-terminal region in ApuRs was found to be highly variable, up to ca. 230 residues in length, with little sequence conservation across the family and no detectable homology to protein domains of known function (Fig. S4b). In multiple ApuR sequences, the C-terminal regions contain clusters of Ser and Thr residues corresponding to clusters of predicted potential phosphorylation sites (see Fig. S4b). Similar clusters in the unstructured C-terminal region (and intracellular loop 3) of G protein-coupled receptors, including opsins, serve as phosphorylation sites involved in triggering desensitization ^67,68^. Phosphorylation at Ser/Thr residues in the C-terminal region is also known for *Cr*ChR1 although the phosphorylation sites are more sparse ^69^. We thus hypothesize that such Ser/Thr clusters might be involved in phosphorylation-mediated regulation of ApuRs in the native cells.

Neither one of the carboxylate residues of the counterion complex of *Hs*BR are fully conserved in ApuRs (Fig 2b,c and Fig. S5). The *Hs*BR carboxylate D85 in TM3 (E123 in *Cr*ChR2) is present in the eight DXQ ApuRs (D80 in *Sf*ApuR2) and is absent in the four XTQ ApuRs. Asp and Glu at this position are characteristic for proton-pumping rhodopsins and most CCRs and are absent from ACRs ^48,70^. The analogy with the cation-transporting rhodopsins is nevertheless incomplete as the ApuRs with Asp at this position lack its hydrogen-bonding partner T89, except for *Sf*ApuR1_DSQ_ which has a hydrogen-bonding Ser instead. In addition, CCRs also typically feature at least one or more carboxylates at other positions in TM2 and TM3 along the ion conduction pathway ^48,71^ which are absent in ApuRs. The second carboxylate residue of the counterion complex in *Hs*BR at position D212, which is found in the majority of regular microbial rhodopsins, is also present as Asp (D213 in *Sf*ApuR2) in all the DXQ ApuRs. However, XTQ ApuRs demonstrate a division between *Ss*ApuR1 and *Mo*ApuR with Glu and *Pk*ApuR and *Ss*ApuR3 with the highly unusual for this position Ser and Thr, respectively, instead.

One of the most salient features of ApuRs is the fixed Gln residue (Q91 in *Sf*ApuR2) at the *Hs*BR proton donor position D96 (see Fig. 2b), a feature they share with NQ sodium and chloride pumps from bacteria, most cryptophyte ACRs and several ACRs from other families. In contrast to most of the ACRs, but similarly to the NQ pumps, Q91 is stabilized by a hydrogen bond with a Ser residue in TM2, S50 (T46 in *Hs*BR; S64 in KR2 ^72^; S54 in NM-R3 ^73^). While in KR2 the Q-S pair constitutes the cytoplasmic gate for sodium ions ^72^, the role of these residues in chloride pumping in NM-R3 is less clear ^73^.

Among the other conserved residues in ApuRs, the fixed Gln at *Sf*ApuR2 position Q24 (M20 in *Hs*BR) in the middle of TM1 stands out as it is rarely encountered in other rhodopsin families (see Fig 2b and Fig. S5). This position is occupied by hydrophobic residues in most ion-pumping rhodopsins and BCCRs but in other ChRs it is most frequently a small polar residue (Ser, Thr or Cys) which constitutes part of the central gate in both CCRs (*Cr*ChR2 residue S63 ^74^) and ACRs (*Gt*ACR1 residue S43 ^75^). The other positions of the ChR central gate are occupied by small amino acids in ApuRs and no other side chain appears to form direct hydrogen bonds with Q24. Nevertheless, the similarly fixed Thr residue T54 (P63 in *Hs*BR, C87 in *Cr*ChR2) in TM2 is also located opposite the retinal-binding Lys close to Q24. Given the high conservation of these two central residues in ApuRs and their location approximately corresponding to the central gate in *Cr*ChR2, Q24 and T54 might be involved in regulating the ion passage as well.

The ApuR predicted structures feature a large open cavity at the extracellular side (extracellular cavity 1, ECC1) above the retinal binding pocket (RBP), and in most of them it connects the pocket to the bulk solvent via a wide tunnel (Fig. 2a). The connection of RBP with the outside in some models appears narrowed or entirely sealed, in particular in ApuRs with Glu at *Sf*ApuR2 position 194 in TM6 which forms a salt bridge with the conserved Arg residue R77 (R82 in *Hs*BR) in TM3 (Fig. S4c). Another open cavity at the cytoplasmic side, ECC2, is discernible in some of the predicted structures, including *Sf*ApuR2 (see Fig. 2 a,b), but its presence and shape are highly sensitive to side chain orientation of the surrounding residues and are variable among structures predicted for the same protein.

All of the predicted ApuR structures feature an opening (fenestration) between transmembrane helices 5 and 6 (TM5 and TM6) (Fig. S4d). RBP fenestrations, also known from diverse other rhodopsin families, vary in position and shape in ApuRs. In the four related ApuRs *Cd*ApuR, *Kc*ApuR, *Tara*ApuR and *Tt*ApuR, the fenestration is due to a small residue (Gly or Ala) at xanthorhodopsin position G156 ^76^ making it similar to the fenestrations in the proteorhodopsin-xanthorhodopsin superclade which enable binding of carotenoid antennas ^77,78^. In other ApuRs the fenestration is more similar to that of *Cr*ChR2.

### Absorption spectra of ApuRs

To characterize ApuR proteins and examine their UV-visible absorption spectra, all intact ApuRs were heterologously expressed in mammalian cultured COS-1 cells. As an initial screening, we performed hydroxylamine bleach assay to check the expression of ApuRs without purification (Fig. S6). This assay uses hydroxylamine to hydrolyze the Schiff base linkage between the retinal chromophore and protein moiety, which induces a spectral change reflecting the disappearance of the original absorption of rhodopsin at its absorption-maximum wavelength (*λ*_max_) and the appearance of absorption of retinal oxime at ∼360 nm. *Sf*ApuR1_DSQ_, *S*sApuR1_ATQ_, and *Mo*ApuR_MTQ_ (the subscripts after the protein names indicate their motif residues TM3) exhibited a broad positive peak in difference spectra in the 449–485 nm region, suggesting that these ApuRs most strongly absorb blue light (Table 1, Fig. S6a). In contrast, *Cd*ApuR_DAQ_, *Kc*ApuR_DAQ_, *S*sApuR2_DAQ_, *Sf*ApuR2_DAQ_, *Sm*ApuR_DAQ_, *Tt*ApuR_DAQ_, and *Pk*ApuR_CTQ_ showed narrow positive peaks around 390–410 nm with negative peaks at 360 nm (Table 1, Fig. S6b), indicating that they likely have *λ*_max_ in the UV-violet region. No clear spectral changes reflecting bleaching of rhodopsins were observed for *Tara*ApuR_DAQ_ and *S*sApuR3_ITQ_ (Fig. S6c), indicating that these two proteins were not properly expressed in the COS-1 cells. Based on sequential alignment, several residues of TM7 in *S*sApuR3_ITQ_ are likely missing, probably impairing the protein expression.

The absolute absorption spectra of nine of successfully expressed ApuRs were measured with purified proteins (Fig. 3). Their *λ*_max_ values are summarized in Table 1. The *λ*_max_ *Mo*ApuR_MTQ_ could not be determined due to strong bleaching during the purification, possibly due to its instability in solubilized conditions. Nevertheless, the results of the hydroxylamine bleach assay (Fig. S6a) demonstrate that *Mo*ApuR_MTQ_ forms a blue-absorbing pigment. Based on the absorption spectra, ApuRs can be classified into UV-absorbing (*Cd*ApuR_DAQ_, *Kc*ApuR_DAQ_, *S*sApuR2_DAQ_, *Sf*ApuR2_DAQ_, *Sm*ApuR_DAQ_, *Tt*ApuR_DAQ_, *Pk*ApuR_CTQ_) and blue-absorbing (*Sf*ApuR1_DSQ_, *S*sApuR1_ATQ_, and *Mo*ApuR_MTQ_) groups. Absorption spectra of UV-absorbing ApuRs exhibited prominent sharp peaks, which are likely to represent vibrational bands.

**Figure 3.**
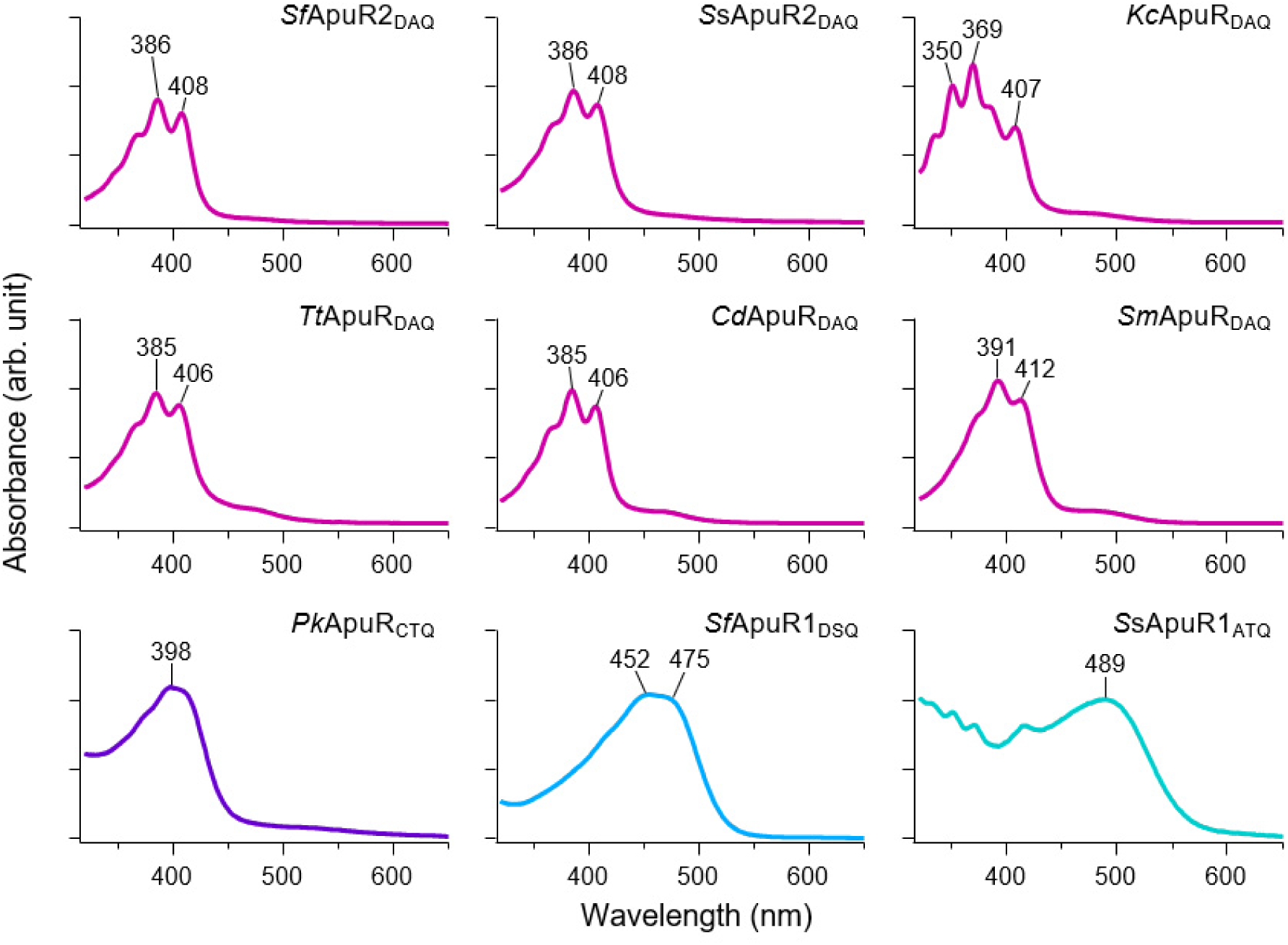
Absorption spectra of ApuRs. Absorption spectra of purified ApuR proteins solubilized in 20 mM HEPES–NaOH, 100 mM NaCl, and 0.05% (w/v) DDM (pH 7.0)

### Ion-transport activity of ApuRs

To evaluate ion transport activity, ApuRs were expressed in ND7/23 cells, and whole-cell patch-clamp measurements were performed. Photocurrents were recorded upon 200-ms illumination at a holding potential ranging from +60 to −80 mV. With standard pipette ([NaCl] = 130 mM) and extracellular ([NaCl] = 145 mM) solutions, nine out of twelve ApuRs exhibited significant photocurrents and inversion from positive to negative currents (Fig. 4a). Despite the similar expression levels in the cells, differences in photocurrent intensity and photocurrent shapes (Figs. 4a and b) suggest a large variation in single-channel conductance and gating dynamics among the ApuRs. Action spectra of ApuRs were estimated by measuring photocurrents under illumination at 377 ± 25, 438 ± 12 and 472 ± 15 nm. The strong photocurrents with 377 ± 25-nm illumination indicates that excitation of deprotonated retinal Schiff base (RSB) forms induces the channel gating in ApuRs (Fig. 4c). Additionally, *Sf*ApuR1_DSQ_, *Mo*ApuR_MTQ_, and *Pk*ApuR_CTQ_, which absorb in the violet and blue regions, exhibited strong photocurrents upon 438 ± 12 and 472 ± 15-nm excitations, indicating that the protonated RSB is responsible for the channel gating in these ApuRs. Three ApuRs, *Tara*ApuR_DAQ_, *S*sApuR1_ATQ_ and *S*sApuR3_ITQ_ exhibited weak photocurrents under UV illumination (Fig. S7). Notably, despite the high membrane localization of *S*sApuR1_ATQ_, its photocurrents were significantly smaller compared to other ApuRs. By contrast, photocurrents were observed in a few cells expressing *S*sApuR3_ITQ_ despite its poor localization to the plasma membrane.

**Figure 4.**
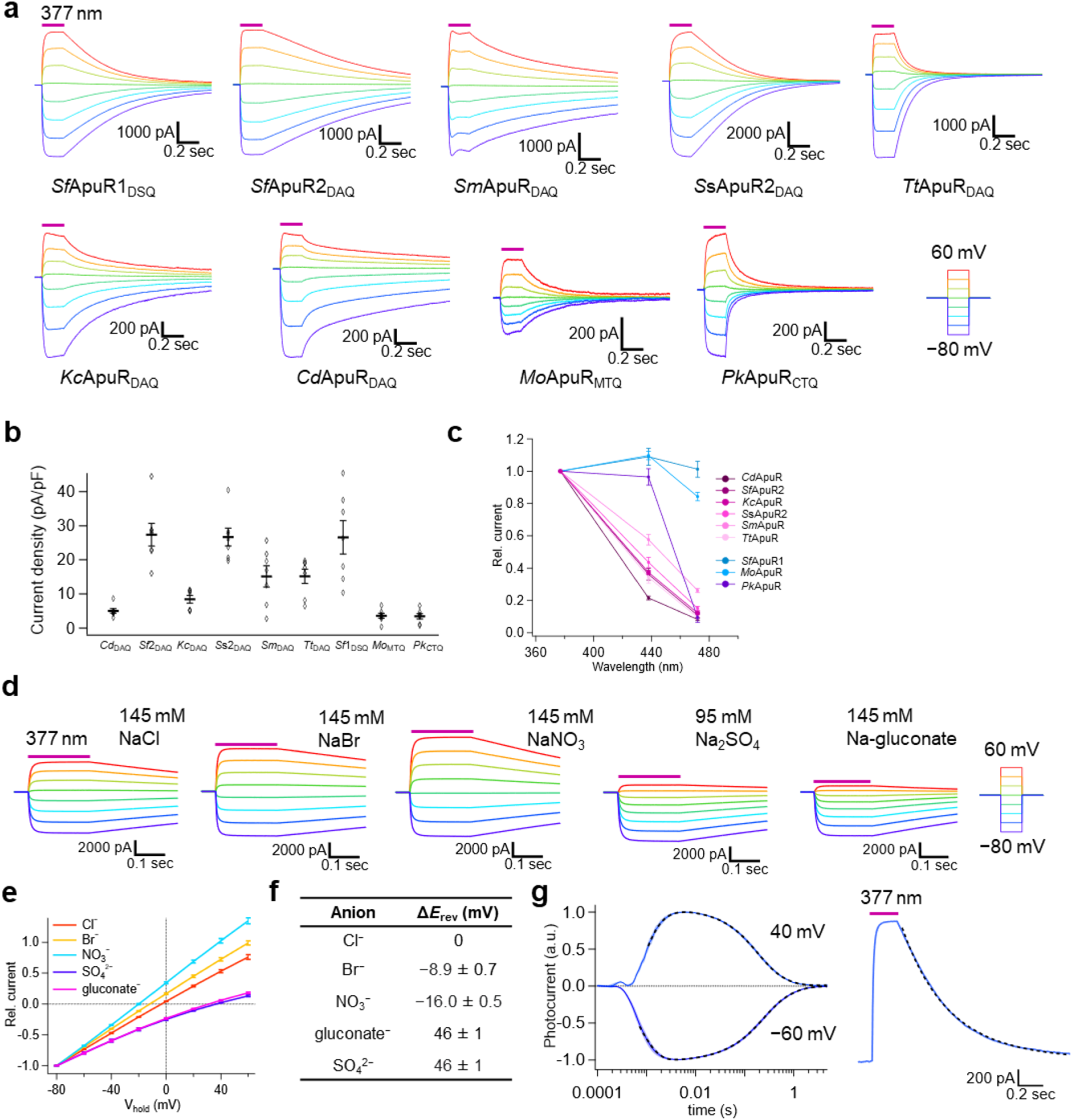
Photocurrents of ApuRs. **a** Photocurrents of ApuRs recorded with standard pipette and extracellular solutions. **b** Steady photocurrents at 40-mV holding potential (mean ± S.E., *n* = 7). Individual data points are shown by white diamonds. **c** Normalized action spectra of ApuRs obtained from photocurrent traces at 40 mV for 200 ms illumination with the light power density of 5.5 mW mm^−2^ (mean ± S.E., *n* = 7). **d** Representative photocurrent traces of *Sf*ApuR2_DAQ_ recorded with extracellular solutions containing different anions. **e** *I–V* curves of the *Sf*ApuR2_DAQ_ steady photocurrent (mean current during the latter half of the illumination time) (mean ± S.E., *n* = 6, photocurrents are normalized at −80 mV.). **f** *E*_rev_ shifts from the extracellular solution containing 145 mM NaCl (mean ± S.E., *n* = 6). **g** Photocurrent traces of *Sf*ApuR2_DAQ_ upon nanosecond laser flash excitation holding potentials of 40 mV and −60 mV recorded with standard pipette and extracellular solutions (left) and representative photocurrent trace of *Sf*ApuR1_DSQ_ recorded at 40 mV upon illumination at 377 nm for 200 ms (right). Fitting curves are shown by broken lines.

To investigate ion selectivity of ApuRs, photocurrents were measured in extracellular solution containing sodium gluconate ([Na-gluconate] = 145 mM). Intensities of positive currents were significantly decreased in *Sf*ApuR2_DAQ_, *Sf*ApuR1_DSQ_, *Sm*ApuR_DAQ_, *Tt*ApuR_DAQ_, *Pk*ApuR_CTQ_ (Figs. 4d, S8a, S8b), which suggests anion transport by ApuRs. Next, photocurrents of *Sf*ApuR2_DAQ_ were measured in extracellular solutions containing different anions (Fig. 4d), and current–voltage (*I*–*V*) curves were plotted in Fig. 4e. The reversal potential (*E*_rev_) was almost zero at 145 mM extracellular Cl^−^. It was shifted to the negative side at 145 mM Br^−^ or NO_3_^−^, while shifting to the positive one at 145 mM gluconate or 95 mM SO_4_^2−^ (Fig. 4f). Among these anions, *Sf*ApuR2_DAQ_ exhibited the highest permeability to NO_3_^−^, similar to ACRs ^79,80^, while showing the lowest permeability to divalent SO_4_^2−^ and larger gluconate.

Regarding the origin of the outward current observed when the external solution contains gluconate or sulfate, two possibilities can be considered: it may reflect the permeation of gluconate or sulfate themselves, or the permeation of protons or hydroxide. To evaluate the proton or hydroxide conductance of *Sf*ApuR2_DAQ_, we analyzed the *I*–*V* curves of *Sf*ApuR2_DAQ_ by changing pH in the extracellular solution of sodium gluconate, sodium sulfate and sodium chloride (Figs. S8d–f). Replacing extracellular solution at pH 7.4 with pH 9.0 and 6.0 resulted in *E*_rev_ shifts (Fig. S8g), indicating that proton/hydroxide permeates *Sf*ApuR2_DAQ_ in the inverse/same direction as chloride, respectively. In NaCl external solutions, the *E*_rev_ shift (Δ*E*_rev_) is smaller compared to other conditions. This is due to the lower permeation of protons or hydroxide ions, which is masked by the stronger permeation of chloride ions. Photocurrent measurements of ApuRs were conducted in an external solution containing 145 mM sodium gluconate (Fig. S8a, b). The positive current, which indicates permeability to proton or hydroxide, differs among ApuRs (Fig. S8c). *E*_rev_ of *Sf*ApuR1_DSQ_ and *Sm*ApuR_DAQ_ are close to that of *Sf*ApuR2_DAQ_ (43 mV with with 145 mM Na-gluconate of extracellular solution at pH 7.4), while those of *Tt*ApuR_DAQ_ and *Pk*ApuR_CTQ_ are close to *Gt*ACR1.Since the best-characterized cryptophyte anion channelrhodopsin, *Gt*ACR1, does not exhibit proton/hydroxide permeability, and its *E*_rev_ is estimated to be 78 mV—closer to the theoretical Cl^−^ Nernst potential (74 mV) than that of *Sf*ApuR2_DAQ_—hence, the positive currents of *Sf*ApuR2_DAQ_ in Na-gluconate and Na_2_SO_4_ extracellular solutions were attributed to proton/hydroxide permeation based on the pH-dependent *E*_rev_ shift.

### Photochemical reaction of SfApuR2_DAQ_ and SfApuR1_DSQ_

To study the photocycle of ApuRs, laser flash photolysis was performed on two representative ApuRs: *Sf*ApuR2_DAQ_ and *Sf*ApuR1_DSQ_, two closely related proteins which differ in their absorption spectra, with absorption maxima in UV (*λ*_max_ = 386 nm) and visible blue light (*λ*_max_ = 452 nm), respectively (Fig. 3).

Fig. 5 shows the transient absorption change of *Sf*ApuR2_DAQ_ excited at 355 nm and *Sf*ApuR1_DSQ_ excited at 450 nm. The photoexcitation of *Sf*ApuR2_DAQ_ resulted in the production of a blue-shifted species in the early step of the photocycle (Figs. 5a, e, and g). This product was assigned to M state and the formation of subsequent visible-absorbing-protonated state (assigned to N state) was observed at 467 nm. Twin-peaked rhodopsins (TwRs) are also known to show the similar photocycle by excitation of UV-absorbing deprotonated state ^81^. UV excitation of deprotonated TAT_LA_, a TwR with TAT motif in TM3, results in the production of a long-lived and visible-absorbing photointermediate with protonated RSB. Compared to TAT_LA_, photoreaction of *Sf*ApuR2_DAQ_ is different, with the absorption change in the UV region occurring prior to the protonation of the RSB. The detailed time evolution of the transient absorption change of *Sf*ApuR2_DAQ_ was recorded using a photomultiplier tube (Fig. 5c). In the early stage of the photocycle, absorption at 386 nm significantly decreased (time constant *τ* = 2.6 μs). In this time region, we were not able to obtain a clear absorption spectrum due to the time resolution of our ICCD linear array detector. Hence, we assume that the M_1_ state is produced prior to the M_2_ state, which was detected by the ICCD linear array detector (Fig. 5g). The time constants of double-exponential N decay were determined to be 250 and 970 ms. The formation and decay of the N state occur near the timing of channel opening and closing (Figs. 4g, 5c and S9), but they do not precisely coincide (Fig. S9a), and the contributions of fast and slow components differ between the absorption and current changes (Figs. S9b and c). Therefore, while the N state corresponds to the open state, the structural changes associated with channel gating are not closely synchronized with the structural changes affecting the retinal absorption. In *Gt*ACR1, the K or L photointermediates represent the ion-conducting state and the formation of the M intermediate corresponds to fast channel closing and its decay to slow channel closing ^82,83^. The relationship between the protonation state of the RSB in the corresponding photointermediate and open/close state of the channel is thus similar between *Sf*ApuR2_DAQ_ and *Gt*ACR1.

**Figure 5.**
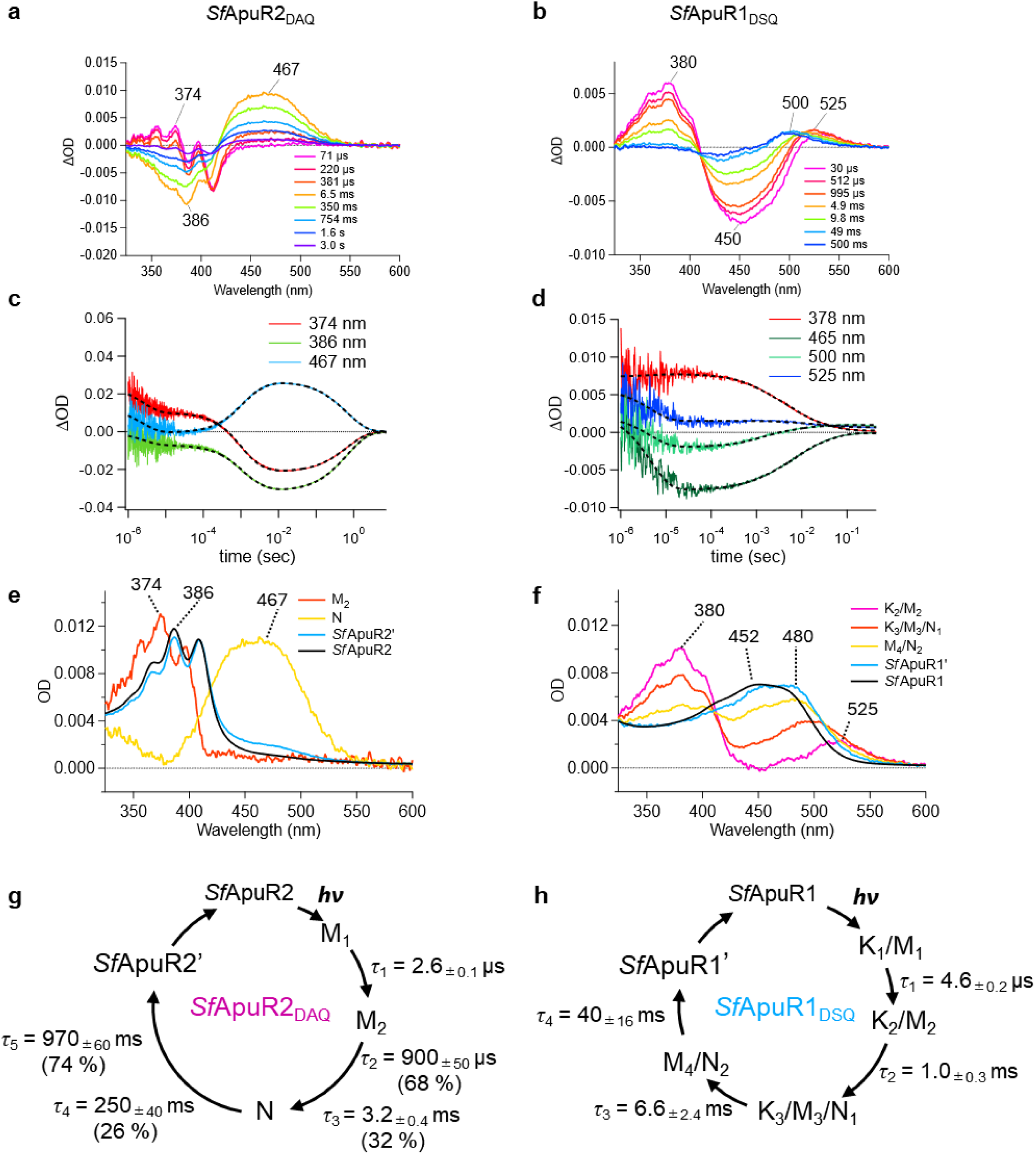
Photocycle of *Sf*ApuR2_DAQ_ and *Sf*ApuR1_DSQ_. **a**, **b** Difference transient absorption spectra of *Sf*ApuR2_DAQ_ (**a**) and *Sf*ApuR1_DSQ_ (**b**). **c**, **d** Time courses of absorption changes of *Sf*ApuR2_DAQ_ (**c**) and *Sf*ApuR1_DSQ_ (**d**). The multi-exponential fitting curves were shown as dashed lines. **e**, **f** Absorption spectra of photointermediates calculated from decay-associated spectra according to Chizhov et al. 1996 ^127^. **g**, **h** The photocycle models of *Sf*ApuR2_DAQ_ (**g**) and *Sf*ApuR1_DSQ_ (**h**).

For *Sf*ApuR1_DSQ_ (Figs. 5b, d, f, and h), the deprotonated M state forms within a time region faster than the time resolution of our measurements, followed by excitation at 450 nm. In addition to the M state, red-shifted absorption was observed at 525 nm (Figs. 5b and f), which was assigned to the K_2_ state. We hypothesize the existence of a primary K_1_ state preceding K_2_ as observed for most microbial rhodopsins ^84^, based on the observation of a significant absorption change at 465, 500 and 525 with a *τ* of 4.6 μs prior to the K_2_ state (Fig. 5d). From the equilibrium state between the K_2_ and M, the absorption in the visible region exhibits a blue shift with a time constant of *τ* = 1.0 ms (Figs. 5b and f). This state was assigned to an equilibrium state between K_3_, M_3_ and N_1_ states which converts to the M_4_/N_2_ state with *τ* = 6.6 ms. Thus, the observed time evolution of absorption changes consists of four components (*τ*_1_ = 4.6 μs, *τ*_2_ = 1.0 ms, *τ*_3_ = 6.6 ms, and *τ*_4_ = 40 ms), none of which match the decay time constants of the photocurrent (*τ*_1_ = 370 ± 40 ms, *τ*_2_ = 1.4 ± 0.2 s, *n* = 7). Therefore, similar to the case of *Sf*ApuR2_DAQ_, the channel gating of *Sf*ApuR1_DSQ_ is likely to be controlled by spectrally silent structural changes of the protein, not synchronized with the structural changes around the retinal. However, considering the channel closing rate, the N state likely corresponds to the open state of the channel as in *Sf*ApuR2_DAQ_.

In both *Sf*ApuR2_DAQ_ and *Sf*ApuR1_DSQ_, the absorption spectra of the latest state calculated from the constant value after the multiexponential transient absorption changes do not precisely match those before excitation (Figs. 5e and f), indicating the existence of long-lived intermediates, *Sf*ApuR2_DAQ_’ and *Sf*ApuR1_DSQ_’, respectively. This result is supported by high-performance liquid chromatography (HPLC) results, which show that the retinal configuration in the dark-adapted state differs from that of the light-adapted state (LA and LA_30 min_ in Figs. S10c).

### Retinal chromophore configurations

The retinal chromophore configurations of *Sf*ApuR2_DAQ_ and *Sf*ApuR1_DSQ_ were analysed using HPLC under varying light conditions (Fig. S10a, b). In the dark-adapted state, the retinal chromophore in *Sf*ApuR1_DAQ_ was predominantly in the all-*trans* form (DA, Fig. S10c, d), which photoisomerized to the 13-*cis* and to a lesser extent to 11-*cis* forms (DA+L). The 11-*cis* component disappeared in the dark within 10 s, resulting in a light-adapted state composed of all-*trans* and 13-*cis* forms (LA, Fig. S10c, d). The subsequent illumination of the light-adapted state induced an increase in the 11-*cis* form and a decrease in the 13-*cis* form (LA+L), leading to isomer proportions similar to those in DA+L. In the dark-adapted state of *Sf*ApuR1_DSQ_, the contribution of the 13-*cis* form was approximately 34%, which is considerably larger than that in *Sf*ApuR2_DAQ_. However, the light-dependent changes in isomer composition of *Sf*ApuR1_DSQ_ were similar to those of *Sf*ApuR2_DAQ_ (Fig. S10c, d). After the cessation of illumination, the 11-*cis* form decreased as in *Sf*ApuR1_DSQ_ (LA_30 min_, Fig. S10c). Taken together, this indicates that photoisomerization occurs between the all-*trans* and 13-*cis* forms and between the all-*trans* and 11-*cis* forms. Additionally, both ApuRs exhibit a light-adapted state that possesses 13-*cis*-retinal. Such a light-adaptation process involving 13-*cis*-retinal is also observed in *Anabaena* sensory rhodopsin from *Anabaena* (Nostoc) sp. PCC7120, the inward H^+^-pumping xenorhodopsin, *Po*XeR, from *Parvularcula oceani*, the channelrhodopsin *Cr*ChR2 and other rhodopsins ^85–87^. Considering this analogy with the photocycles of *Po*XeR and *Cr*ChR2 which demonstrate branching reactions ^86^, we propose a photoisomerization model of the retinal in *Sf*ApuRs with an additional photocycle involving 11-*cis*-retinal as in Fig. S10e. While the all-*trans*-to-13-*cis* photoisomerization is likely to lead the channel opening as in *Cr*ChR2 and *Gt*ACR1 ^88^, it is unclear how 11-*cis*-retinal contributes to the function of ApuRs.

### Spectral-tuning sites

To get insights into the potential mechanism responsible for the switching between absorption in UV and in blue among ApuRs, we correlated the critical residues around the RSB with the absorption spectra of the proteins. First, we noticed that the blue-absorbing ApuRs are included in both groups of ApuRs: in the DXQ ApuRs, which have two Asp residues in the counterion complex (D80 and D213 in *Sf*ApuR2_DAQ_), and in the XTQ ApuRs, which either have one Glu in TM7 or have no carboxylates in the vicinity of the Schiff base (see *Structural features* and Table 1). This division implies that the color-switching mechanism is different in the two groups. The only DXQ ApuR to absorb in blue is *Sf*ApuR1_DSQ_ which exceptionally has a Ser residue forming a hydrogen bond with the Asp residue in TM3 instead of Ala of the UV-absorbing DAQ ApuRs. We thus hypothesized that the second site of the motif plays a central role in the spectral tuning in DXQ ApuRs. To test this hypothesis, we examined the absorption spectra of mutants of *Sf*ApuR1_DSQ_ and *Sf*ApuR2_DAQ_ in which the residue in the second site of the motif was reciprocally swapped: *Sf*ApuR2_DAQ_ A84S (DSQ-mutant) and *Sf*ApuR1_DSQ_ S73A (DAQ-mutant). *Sf*ApuR2_DSQ_ A84S exhibited a substantial absorption peak in the visible region (450–478 nm), while the decreased peaks in the UV regions were still observed (Fig. 6a). In contrast, the *Sf*ApuR1_DAQ_ S73A has the largest absorption peak in the UV region with a weaker peak in the visible region (Fig. 6b). Therefore, the Ser and Ala residues at the second position of the motif is essential for *Sf*ApuR2 and SfApuR1 to absorb visible and UV light, respectively, suggesting that the second site of the motif acts as a switch for the protonation state of the RSB in ApuRs with two Asp residues near the RSB. Interestingly, the action spectra of *Sf*ApuR2_DSQ_ A84S and *Sf*ApuR1_DAQ_ S73A mutants showed increases in the photocurrent upon visible and UV excitation, respectively, compared to the WT proteins (Fig. 6d and e). These shifts in the action spectra are consistent with the absorption spectra, indicating both the protonated and deprotonated states in the same protein can contribute to the ion transport, enabling ApuRs to alter their absorption without impairing channeling function.

**Figure 6.**
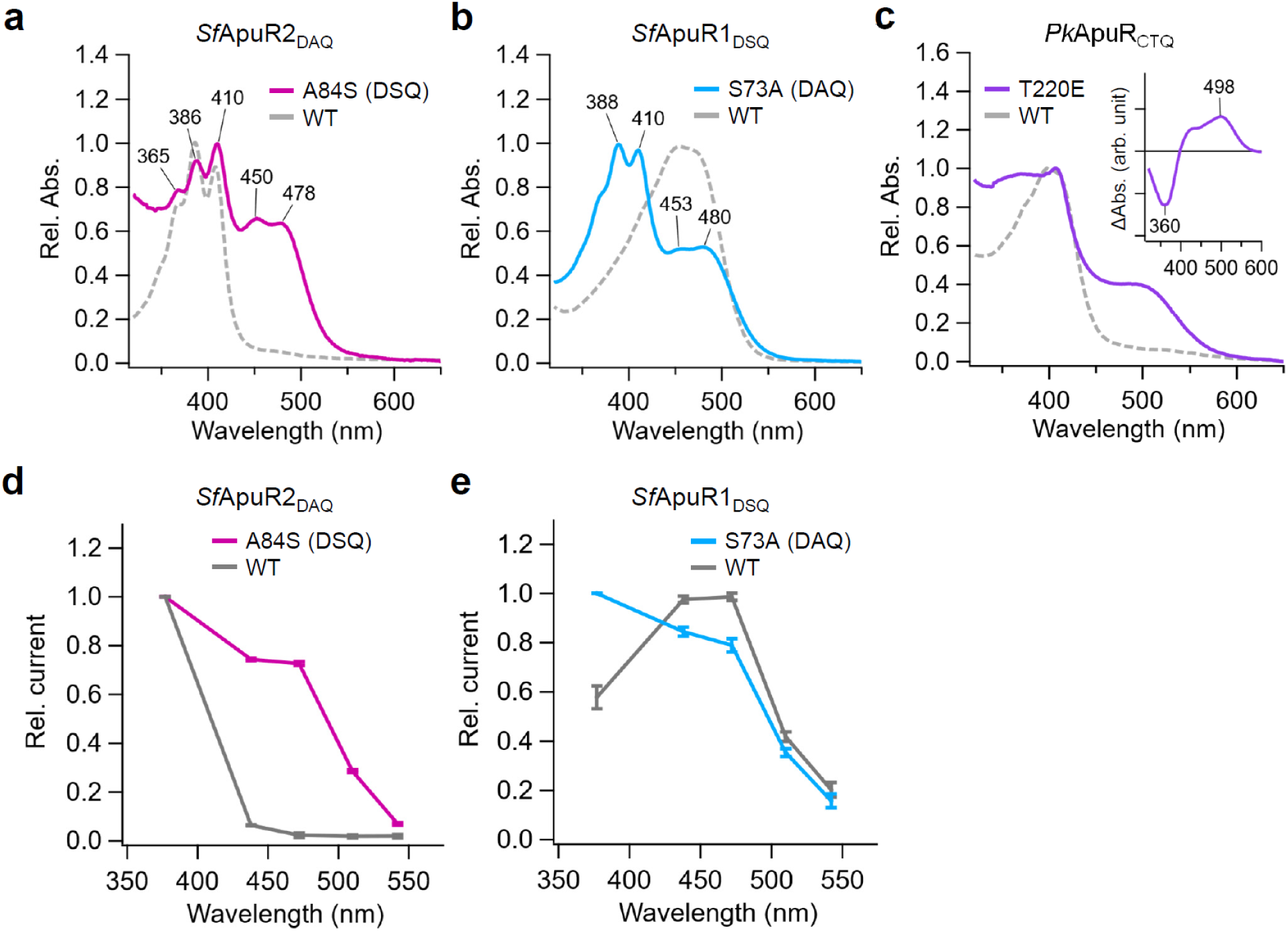
Spectral-tuning sites of three ApuRs. **a**-**c** Absorption spectra of *Sf*ApuR2_DAQ_ A84S (**a**), *Sf*ApuR1_DSQ_ S73A (**b**), and *Pk*ApuR_CTQ_ T220E (**c**) mutants are shown with those of WT. (**c**, inset) A difference spectrum upon hydroxylamine bleach of the purified *Pk*ApuR_CTQ_ T220E. **d**, **e** Normalized action spectra of *Sf*ApuR2_DAQ_ A84S (**d**) and *Sf*ApuR1_DSQ_ S73A (**e**) obtained from photocurrents at −40 mV with 2-ms illumination with the light power density of 0.5 mW mm^−2^ (mean ± S.E., *n* = 5). The motif residues of TM3 were shown in parentheses.

We reasoned that in the case of the XTQ ApuRs, the defining trait underlying the division between the UV- and blue-absorbing forms is the presence of Glu in TM7, since *Pk*ApuR_CTQ_ with the unusual Thr residue at this position and thus lacking any potential counterions is the only successfully expressed XTQ ApuR to absorb in UV (Tabel 1 and Figs. 3 and S6). Therefore, we examined the absorption spectra of a T220E mutant of *Pk*ApuR_CTQ_ to explore the mechanism underlying its UV absorption. As a result, *Pk*ApuR_CTQ_ T220E exhibited an additional absorption peak in the blue region (Fig. 6c) thus emulating the phenotype of the natural ApuR variants with Glu at this position. This indicates that the RSB is deprotonated in the *Pk*ApuR_CTQ_ WT and protonated at least partly in this mutant, as the Glu introduced at position 220 acted as a counterion.

### Mutational analysis of closing kinetics in SfApuR2_DAQ_

We explored key residues for the gating mechanism of *Sf*ApuR2_DAQ_. From the comparison of the amino acid sequences of *Sf*ApuR2_DAQ_, *Hs*BR, and channelrhodopsins, we focused on the following residues: A61, T85, C106, and C216 (Fig. 7a with the side chains from of *Hs*BR ^89^ and S5). The A61 residue in TM2 corresponds to a highly conserved Tyr in *Hs*BR (Y57) and many other microbial rhodopsins. The photocurrents of A61Y in *Sf*ApuR2_DAQ_ exhibit a significant deceleration in the closing rate (Figs. 7b and c). At C216, one position before the retinal-binding K217 in TM7, while some microbial rhodopsins such as *Gt*ACR1 and *Gt*ACR2 have Cys, *Hs*BR and many other microbial rhodopsins have Ala, Ser, Thr, and Asn ^90^. Similarly to A61Y, *Sf*ApuR2_DAQ_ C216A showed a significant slowdown in the closing rate (Fig 7b). Probably due to the highly prolonged conducting state of C216A, the cell membrane became leaky during the illumination, preventing the determination of the closing rate. At T85 position in TM3, while most channelrhodopsins have Cys (e.g., C128 in *Cr*ChR2 forming the so-called DC-gate with D156 in TM4), and the mutation of this residue slows down the gating rates ^91^, interestingly, *Sf*ApuR2_DAQ_ T85C exhibited acceleration of the closing rate (Figs. 7b and c). *Sf*ApuR2_DAQ_ C106L, a mutant mimicking *Hs*BR L111 in TM4, exhibited double-exponential closing dynamics in which components faster and slower than the closing of the WT were observed (Figs. 7b, c). For *Sf*ApuR2_DAQ_ T85C and C216A, these positions are close to the retinal, suggesting that the point mutations may directly alter the rate of the photoreaction like *Cr*ChR2 C128T mutant ^92^. In contrast, A61Y and C106L are located farther from the retinal. Particularly, C106 is distal from the chromophore, but this residue in TM4 likely to form hydrogen bonds between side chains and main chains with H93 in TM3, in the AlphaFold structural model, suggesting helix-helix interactions across TM3– TM4. Since helix movements are expected to be crucial for gating, the change in the closing rate in *Sf*ApuR2_DAQ_ C106L may be attributed to weakened inter-helical interactions. In the case of *Sf*ApuR2_DAQ_ A61Y, the with a tyrosine residue may bring the side chain closer to D80 in TM3 and D213 in TM7, putative counterions for protonated RSB. This proximity would provide stronger interactions, potentially stabilizing the N state. Moreover, the enhanced interactions between the candidate counterions and tyrosine could strengthen inter-helical interactions between TM2, TM3, and TM7, potentially slowing down the critical helical movements necessary for channel closing.

**Figure. 7.**
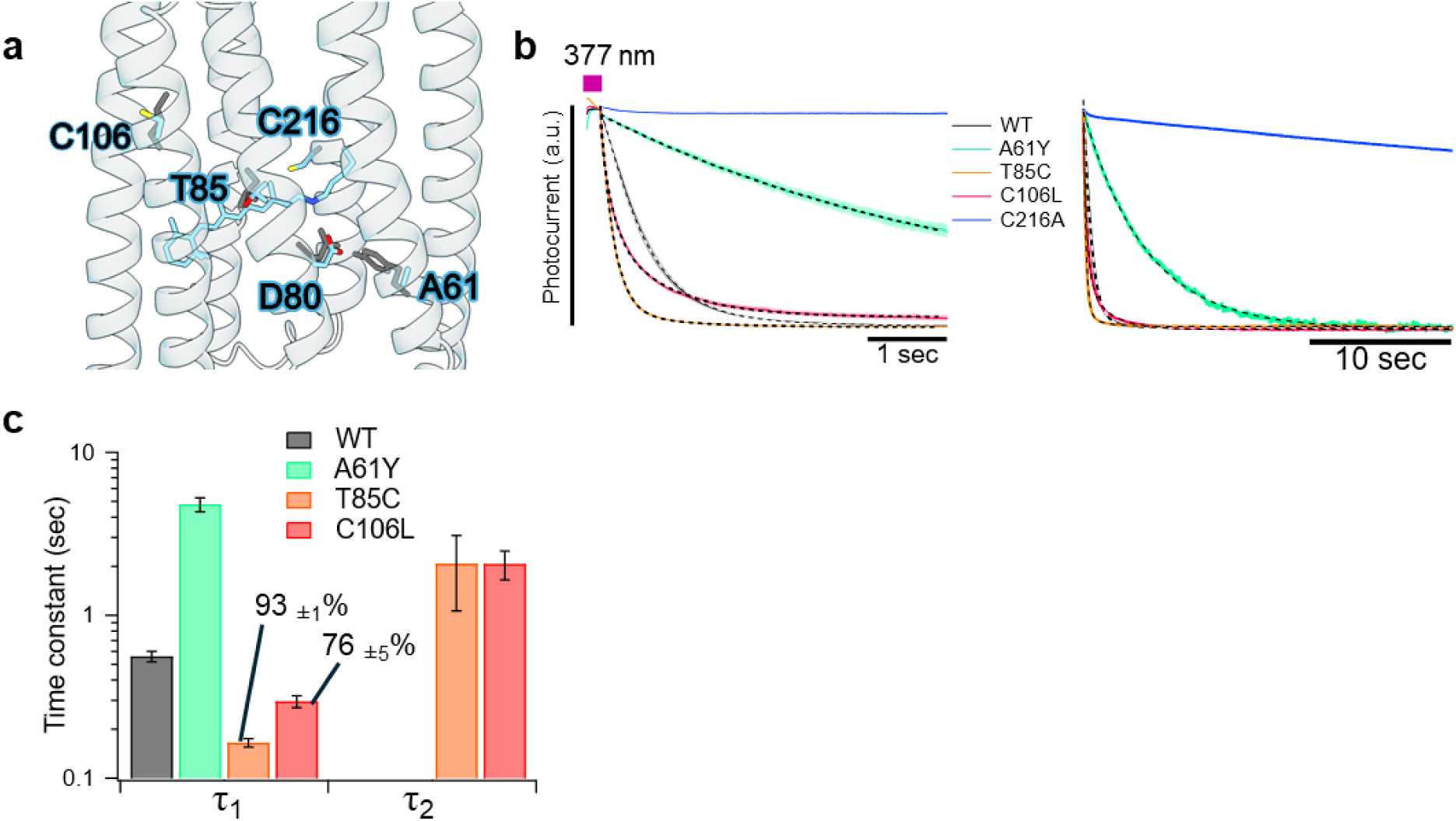
Effect of point mutation on the channel gating of *Sf*ApuR2_DAQ_. **a** AlphaFold3 structural model for *Sf*ApuR2_DAQ_ (blue) showing the mutated residues. The retinal structure and some side chains (gray sticks) are of *Hs*BR (PDB ID: 1KGB) ^89^. **b** Photocurrent decays of *Sf*ApuR2_DAQ_ WT and its mutants after 200-ms illumination recorded in shorter (left) and longer (right) time ranges. Fitting curves are shown as dashed lines. **c** Time constants of channel closing of the *Sf*ApuR2_DAQ_ WT and mutants (mean ± S.E., *n* = 3–5).

## Discussion

### ApuRs, a new family of microbial rhodopsins

Our photocurrent experiments indicate that ApuRs act as light-gated ion channels transporting mainly monovalent anions. Given the lack of phylogenetic ties with the family of channelrhodopsins, ApuRs represent an independent evolution of ion channeling activity among microbial rhodopsins and are in fact structurally more similar to trimeric proton-pumping rhodopsins.

A number of conserved residues inside the proteins indicate that the ion pore runs in ApuRs through the rhodopsin monomer as in channelrhodopsins. Two conserved Gln residues: Q91 in the intracellular half of TM3 and Q24 in the center of TM1 at a position of one of the residues of the central gate in ChRs, are the prime candidates for forming the core of the channel gates in ApuRs. Q91 constitutes the only residue of the TM3 motif universally conserved in ApuRs and in the predicted models it forms a hydrogen bond with the Ser residue at S50.

Despite some parallels in the critical residues between ApuRs and other microbial rhodopsins, they represent convergences and not inherited similarities. Thus, given the lack of direct phylogenetic ties between the NQ pumps and ApuRs, the Q-S pair must have evolved independently in these families. Similarly, the Asp residue at D85 in *Hs*BR, characteristic to proton-pumping rhodopsins and cation ChRs, is a secondary development among ApuRs (see Fig. 1b) unrelated to the ancestral proton pumping activity of microbial rhodopsins ^93^.

### Physiological function of ApuRs

Our results show how ApuRs are widespread across apusomonads, they branch deeply within the diversity of type 1 rhodopsins, and their evolution broadly follows the known apusomonad phylogeny (Fig. 1b). ApuRs’ sequence conservation and diversification suggest that they fulfill an important physiological function in these HFs.

All of the analyzed ApuRs demonstrate a strong tendency for blue-shifted absorption spectra. To the best of our knowledge, the UV-absorbing ApuRs are in fact the most blue-shifted rhodopsin channels. Mutational analysis revealed two switches— corresponding to D80 and D213 in *Sf*ApuR2_DAQ_— which allowed ApuRs to frequently change their absorption between the UV- and blue-absorbing forms in the DXQ and the XTQ ApuRs, respectively. Moreover, the two species which harbor multiple ApuR genes: *Singekia* sp. FB-2015 and *S. franciliensis*, consistently have both blue- and UV-absorbing ApuRs, thus together covering the whole range of short wavelengths. Together this indicates that short blue and UV light play a crucial role in the physiological function of ApuRs. Absorption in the UV-blue range speaks against utilization of carotenoid antennas by ApuRs despite the presence of xanthorhodopsin-like fenestrations in some ApuRs.

Since apusomonads inhabit sediments in soils, freshwater and marine settings ^54,55^, the finding of photoreceptors in them is at first sight surprising. Light absorption of the different ApuRs was found between blue and UV light spectrum. Nevertheless, Shibata *et al.* (2024) ^94^ have recently demonstrated negative phototactic behavior (avoidance) in the ApuR-possessing apusomonad *P. kaiyoae* under blue light. Blue light might help these protists to orient in the surface layer of the sediment and/or avoid photodamage ^94^. In line with this, we have found that the ApuR protein of *P. kaiyoae*, *Pk*ApuR_CTQ_, exhibits the absorption band which peaks at 398 nm and covers the blue-light region where the negative phototaxis is observed (Fig. 3). It is thus tempting to connect ApuRs to the observed avoidance behavior, but direct experiments on apusomonad cells (such as ApuR knock down, drug assays including hydroxylamine treatment and patch clamping) would be required to test this hypothesis.

ApuRs showcase that a new diversity of microbial rhodopsins await discovery within the diversity of heterotrophic flagellates. It is also worth highlighting that we could identify only two ApuR types in environmental datasets ^95^. Apusomonads are protists that inhabit mainly sediments, and these datasets originate from water-column samples, which may explain the relative scarcity of these rhodopsins in environmental data. Thus, our study showcases how using culture-based methods can be advantageous when it comes to exploring the diversity of rhodopsins in protists poorly represented in environmental surveys.

Our electrophysiological measurements clarified the light-gated anion-selective ion channel function of ApuRs. Interestingly, both UV- and violet/blue-absorbing forms exhibit similar channel activity, indicating that the protonation state of the RSB in the dark state while playing a critical role in the color tuning is not important for the channel function. The closing rate of *Sf*ApuR2_DAQ_ is regulated by several residues whose mutations cause significant changes in the closing kinetics. Particularly, mutational experiments identified that the difference in the second motif residue between alanine and serine in DXQ ApuRs is critical to the regulation of the shift between the UV and blue absorption, depending on the protonation state of the RSB. *Sf*ApuR2_DAQ_ C216A, showing drastically extended open state, can be used as a highly sensitive optogenetics tool that inhibits neural activity with a small amount of illumination. For *Sf*ApuR2_DAQ_ and *Sf*ApuR1_DSQ_, the N state is likely to represent the open state, indicating that the protonation of the RSB is essential for the channel opening. In order to understand the gating and color-tuning mechanisms, the proton transfer pathway in ApuRs will be investigated by 3-D structural analysis and spectroscopic measurements such as time-resolved infrared and Raman spectroscopy in the near future.

## Methods

### Collection and searches of ApuRs

To study the presence or absence of microbial rhodopsins in apusomonads blastp, tblastn ^96^ and custom HMM profiles ^97^ searches were performed against all available proteomes of apusomonads ^57,98–101^. We also performed searches against GenBank non-redundant databases ^96,102^ (last searched in January 2024) and EukProt v. 3 ^99^. Environmental searches were performed on the Marine Atlas of Tara Oceans Unigenes (MATOU) ^62^, Marine Microbial Eukaryote Transcriptome Sequencing Project (MMETSP) ^103,104^, multiple assemblies of the Tara Oceans prokaryotic/girus metagenomic data ^105,106^. In all cases we used the proteins ApuRs from *P. kaiyoae* and *T. trahens* as queries.

### ApuR gene phylogeny

The phylogenetic dataset is a modified version of the dataset by Rozenberg et al. 2021^9^, including the ApuR sequences for a total of 396 sequences. The sequences were aligned with MAFFT v.7 ^107^ and trimmed with trimAL v.1.2 ^108^ using -gt 0.3 (to remove sites with gaps in greater than 70% of the sequences) and manual inspection. The trimmed alignment was taken to maximum likelihood phylogenetic reconstruction with IQ-TREE v.1.6.11 ^109^ under the Q.pfam+F+R8 model as chosen per BIC. Statistical support was generated with 1,000 ultrafast bootstraps. Sequences, phylogenetic dataset and trees are deposited on figshare (https://doi.org/10.6084/m9.figshare.26965591.v2). Gene tree-species tree reconciliation was performed with Notung v. 2.9.1 ^110^ using the default costs.

### Structural analyses

Structures of ApuR rhodopsin domains were predicted using the AlphaFold 3 package ^111^ specifying retinal as the covalent modification. The default set of the databases was used, with the addition of a custom database of rhodopsin sequences extracted from EukProt v. 3.0 ^99^, Global Microbial Gene Catalog v.1.0 ^112^, Tara Oceans Eukaryote Gene Catalog v.1 ^62^, Marine Eukaryotic Reference Catalog ^64^, MGnify clusters v. 2023_02 ^113^ and Ocean Microbial Reference Catalog v. 2 ^114^, supplied to increase the space of sequences recruited for alignment. Twenty seeds were used with five samples per seed and the best-ranking model was generally picked per protein. For some proteins, we found it necessary to filter out models with an unusual orientation of the side chain of tyrosine at *Hs*BR position Y185. The structures were minimized with GROMACS v. 4.5.7 ^115,116^ as implemented in the preparatory steps of the PyARM workflow ^117^. For analysis of oligomeric structures, di-, tri- and pentamers were predicted for the ApuRs and reference rhodopsins using truncated versions of the proteins in a similar way as for the monomers, except providing five seeds.

Sequence conservation was analyzed using the ConSurf server ^118^. Pairwise structural alignments to rhodopsin structures in Protein Data Bank were performed with TM-align v. 20170708 ^119^. Cavities were analyzed with PyVOL v. 1.7.8 ^120^. Structure manipulations and visualization were performed using the application programming interface and graphical user interface of PyMOL v. 2.6.0 ^120^.

Prediction of potential phosphorylation sites in the C-terminal extensions of ApuRs and reference rhodopsins was performed with NetPhos v. 3.1 ^121^. Only “non-specific” sites passing the score threshold of 0.75 were included.

### DNA constructs for protein expression

DNA fragment of each apusomonad rhodopsin was synthesized after the codon-optimization for expression in human cells and inserted into pUC57 vector (GenScript, NJ). For the hydroxylamine bleach assay and electrophysiology, ApuRs were subcloned into p*Gt*CCR4-3.0-eYFP ^122^ by replacing *Gt*CCR4 with ApuRs using the restriction enzymes, EcoRI and BamHI. In these constructs, ApuRs were fused with two protein trafficking signals and eYFP at their C-termini. ApuRs were tagged with 6×His tag at their C-termini and inserted into pcDNA3.1 for protein expression in mammalian cultured cells for purification. ApuRs *Sf*ApuR1_DSQ_ and *Sf*ApuR2_DAQ_ were inserted into the expression vector pPICZA for protein expression in *Pichia pastoris* cells. *Sf*ApuR1_DSQ_ was tagged with 6×His tag at the C-terminus. *Sf*ApuR2_DAQ_ was fused with a Kir2.1 membrane trafficking signal, followed by HRV3C site, EGFP as an expression marker, and 8×His tag. Point mutations were introduced by the QuikChange^®^ site-directed mutagenesis kit (Agilent Technologies, Santa Clara, CA).

### Protein expression and purification with COS-1 cells for UV-visible spectroscopy

Cell culture and transfection were performed as described elsewhere ^123^. Briefly, COS-1 cells were cultured using Dulbecco’s modified Eagle’s medium (D-MEM, FUJIFILM Wako Pure Chemical Co., Japan) supplemented with 10% fetal bovine serum (FBS) and transfected with polyethylenimine ^124^. To reconstitute rhodopsin, cells were supplemented with all-*trans*-retinal (ATR; 2.5 µM, final conc.) on the following day and harvested 48 hrs after transfection.

For hydroxylamine bleach assay, the harvested cells were solubilized with 3% (w/v) *n*-dodecyl-β-D-maltoside (DDM) in a phosphate buffer (pH 7.0) containing 53 mM phosphate and 106 mM NaCl. After centrifugation (5,000 ×*g*, 20 °C, 5 min), the supernatant was mixed with 2 M hydroxylamine (pH 7.0; final concentration: 50 mM). Absorption spectra were repeatedly measured after different periods of time using a UV-visible spectrophotometer (V-750, JASCO, Japan).

For protein purification, the harvested cells were rinsed with a buffer containing 20 mM 4-(2-hydroxyethyl)-1-piperazineethanesulfonic acid (HEPES)–NaOH (pH 6.5), 140 mM NaCl, and 5 mM MgCl_2_ and sonicated by an ultrasonic homogenizer (VP-300N, TAITEC, Japan) in a buffer containing 50 mM Tris(hydroxymethyl)aminomethane (Tris)– HCl (pH 8.0) and 300 mM NaCl (buffer A). The membrane fraction was collected by ultracentrifugation (142,000 ×*g*, 4 °C, 1 h) and solubilized with a buffer A containing 1% (w/v) DDM. The supernatant was collected after centrifugation (5,000 ×*g*, 20 °C, 5 min). Purification was conducted using TALON spin columns (Takara Bio, Japan) according to the manufacturer’s instructions. The resin was washed with a buffer A containing 0.1% (w/v) DDM and 5 mM imidazole. Elution was performed with a buffer containing 50 mM Tris (pH 7.5), 300 mM NaCl, 300 mM imidazole with 0.05% (w/v) DDM. The buffer was exchanged to a buffer containing 20 mM HEPES–NaOH (pH 7.0), 100 mM NaCl, and 0.05% (w/v) DDM using an ultrafiltration spin column.

### Protein expression and purification with P. pastoris for transient absorption measurements

*Sf*ApuR1_DSQ_ tagged with 6×His and *Sf*ApuR2_DAQ_ tagged with EGFP followed by 8×His were expressed in *P*. *pastoris* SMD1168H strain. Pre-culture was carried out using BMGY medium for 2 days at 30 °C. The cells were harvested 2 days after the protein expression was induced by BMMY medium (0.5% (v/v) methanol) containing 30 μM ATR at 30 °C. The cultured cells were collected by centrifugation and resuspended in 50 mM Tris–HCl, 7 mM dithiothreitol (DTT), 25 μM ATR, 0.1% (w/v) zymolyase (Nacalai tesque, Japan), protease inhibitor (cOmplete^TM^, Roche, Switzerland), and a small amount of DNase at pH 7.5. The cell suspensions were slowly shaken in the dark at room temperature for 3–4 hrs. Then, the suspensions were vortexed more than 10 times with a 50% volume of zirconia beads (*Φ* = 0.5 mm) for 1 min, and centrifuged at 700 ×*g*. The supernatants were centrifuged for 20 min at 185,300 x*g*, and membrane fraction was collected, then the membrane fraction was homogenized and resuspended in a solubilization buffer (20 mM Tris–HCl (pH 7.5), 150 mM NaCl, 10% (v/v) glycerol, 2% (w/v) DDM, 1 mM phenylmethylsulfonyl fluoride (PMSF)) at 4 °C for 1 h. The suspensions were centrifuged for 20 min at 185,300 x*g*, then the supernatant was loaded in a HisTrap^TM^ HP column (Cytiva, MA). The column was washed with an equilibration buffer containing 20 mM HEPES–NaOH (pH 7.0), 150 mM NaCl, 10% (v/v) glycerol, 0.05% (w/v) DDM, and then treated with an elution buffer containing 50 mM Tris–HCl (pH 7.5), 300 mM NaCl, 500 mM imidazole, 0.1% (w/v) DDM. The eluted *Sf*ApuR2_DAQ_ and *Sf*ApuR1_DSQ_ and were dialyzed against the equilibration buffer at 4 °C. For *Sf*ApuR2_DAQ_, fused EGFP followed by 8×His was digested by turbo3C protease (Nacalai tesque, Japan) during dialysis. The solution of *Sf*ApuR2_DAQ_ was applied to a HisTrap^TM^ HP column again, and the digested sample was eluted. The eluted sample was dialyzed with the equilibration buffer.

### Cell culture and transfection for electrophysiology

ND7/23 cells were grown in D-MEM supplemented with 5% FBS under a 5% CO_2_ atmosphere at 37 °C. ND7/23 cells were attached onto collagen-coated 12-mm coverslips (IWAKI, cat. 4912-010, Japan) placed in a 4-well cell culture plate (SPL Life Sciences, cat. 30004, Korea). The expression plasmids were transiently transfected in ND7/23 cells using Lipofectamine^®^ 3000 transfection reagent (Thermo Fisher Scientific Inc., MA). Seven to eight hours after the transfection, the medium was replaced with D-MEM containing 10% horse serum (New Zealand origin, Thermo Fisher Scientific Inc., MA), 50 ng mL^−1^ nerve growth factor-7S (Sigma-Aldrich, MO), 1 mM N^6^,2’-O-dibutyryladenosine-3’,5’-cyclic monophosphate sodium salt (Nacalai tesque, Japan), 1 μM cytosine-1-β-D(+)-arabinofuranoside (FUJIFILM Wako Pure Chemical Co., Japan), and 2.5 μM ATR. Electrophysiological recordings were conducted at 1–3 days after the transfection. The transfected cells were identified by observing the eYFP fluorescence under an up-right microscope (BX50WI, Olympus, Japan).

### Electrophysiology

All experiments were carried out at room temperature (20–22°C). Currents were recorded using an EPC-8 amplifier (HEKA Electronic, Germany) under a whole-cell patch clamp configuration. The data were filtered at 1 kHz, sampled at 50 kHz (Digidata1440 A/D, Molecular Devices Co., CA) and stored in a computer (pClamp11.1, Molecular Devices Co., CA). The standard internal pipette solution for the whole-cell voltage clamp recordings from the ND7/23 cells contained (in mM) 130 NaCl, 5 Na_2_EGTA, 1 MgCl_2_, 10 HEPES, 2.5 MgATP, 0.0025 ATR (pH 7.4 adjusted with NaOH). The standard extracellular solution contained (in mM): 145 NaCl, 2.5 CaCl_2_, 1 MgCl_2_, 10 HEPES, and 11 glucose (pH 7.4 adjusted with NaOH). In the ion selectivity measurement, the extracellular solution contained (in mM) 145 NaX (X = Cl^−^, Br^−^, NO_3_^−^, gluconate^−^), 2.5 CaCl_2_, 1 MgCl_2_, 10 HEPES, 11 glucose (pH 7.4 adjusted with NaOH). For the divalent anion selectivity measurement, the extracellular solution contained (in mM) 95 Na_2_SO_4_, 2.5 CaCl_2_, 1 MgCl_2_, 10 HEPES, 11 glucose (pH 7.4 adjusted with NaOH). The liquid junction potentials (LJPs) were calculated by pClamp 11.1 software (Table S1) and estimated reversal potentials were compensated by calculated LJPs. For the measurement of pH dependence of *I–V* curve, components of extracellular solutions were based on the 145 NaX (X = Cl^−^, gluconate^−^) and Na_2_SO_4_ solutions described above, in which 10 mM HEPES was exchanged for 2-morpholinoethanesulfonic acid (MES) at pH 6.0, or for *N*-cyclohexyl-2-aminoethanesulfonic acid (CHES) at pH 9.0.

For whole-cell voltage clamp, illumination at 377 ± 25, 438 ± 12, 472 ± 15, 510 ± 5, and 542 ± 13 nm were carried out using a SpectraX light engine (Lumencor Inc., OR) controlled by pClamp 11.1 software. ApuRs were illuminated through an objective lens (LUMPlan FL 40x, NA 0.80W, Olympus, Japan). For the standard photocurrent detection, the power of the 377-nm light was directly measured under a microscope using a visible light-sensing thermopile (MIR178 101Q, SSC Co., Ltd., Japan) and was adjusted to 5.5 mW mm^−2^. The action spectrum was measured at a holding potential of 40 mV at wavelengths with equivalent photon density of 5.5 mW mm^−2^ for nine ApuRs. For the comparison of the WT and the mutants on the second motif residue of TM3 of *Sf*ApuR2_DAQ_ and *Sf*ApuR1_DSQ_ the action spectra were measured at a holding potential of −40 mV at wavelengths with equivalent photon density of 0.5 mW mm^−2^. For the laser-flash patch clamp experiment, a laser flash (3–5 ns) at 355 nm (Nd:YAG laser, Minilite II, Continuum, CA) was used. In the laser-flash patch clamp experiment, the series resistance was compensated for by 70%.

### High-performance liquid chromatography (HPLC) analysis

The chromophore configuration of ApuRs was analyzed by HPLC as described elsewhere ^125^. Briefly, the chromophore retinal was converted to retinal-oxime by adding methanol and hydroxylamine and extracted with *n*-hexane. The extracts were analyzed using an HPLC system composed of a silica column (particle size 3 μm, 150 × 6.0 mm; Pack SIL, YMC, Japan), a pump (PU-4580, JASCO, Japan), and a UV-visible detector (UV-4570, JASCO; detection, 360 nm) using a *n*-hexane-based solvent mixture (15% ethyl acetate and 0.15% ethanol in *n*-hexane) as the mobile phase.

### Laser flash photolysis

The experimental setup for the laser flash photolysis measurement was similar to that reported previously ^126^. The purified protein samples were solubilized in a solvent containing 20 mM HEPES–NaOH (pH 7.0), 150 mM NaCl, 0.05% (w/v) DDM, 10% (v/v) glycerol. The absorption of *Sf*ApuR2_DAQ_ and *Sf*ApuR1_DSQ_ solution was adjusted to 0.3–0.5 at excitation wavelengths of 355 nm for *Sf*ApuR2_DAQ_ and 450 nm for *Sf*ApuR1_DSQ_. For *Sf*ApuR2_DAQ_, the sample was illuminated with a beam of third harmonics of a nanosecond-pulsed Nd:YAG laser (*λ* = 355 nm, 5.1 mJ cm^−2^, 0.1–0.3 Hz) (INDI40, Spectra-Physics, CA). For the excitation of *Sf*ApuR1_DSQ_ at 450 nm, we used an OPO system (pulse width: ∼10 ns, pulse energy = 5.7 mJ cm^−2^, BasiScan, Spectra-Physics, CA, USA) pumped by the third harmonics of a nanosecond-pulsed Nd:YAG laser (INDI40, Spectra-Physics, CA). The time course of the transient absorption change was obtained by observing the intensity change of the output of a Xe arc lamp (L9289-01, Hamamatsu Photonics, Japan) monochromated by a monochromator (S-10, SOMA OPTICS, Japan) and passed through the sample after photoexcitation using a photomultiplier tube (R10699, Hamamatsu Photonics, Japan) equipped with a notch filter (532 nm, bandwidth = 17 nm) (Semrock, NY) to remove the scattered pump pulse. The signals were globally fitted with a multiexponential function to determine the lifetimes and absorption spectra of each photointermediate.

## Data availability

Sequences, structure predictions, phylogenetic dataset and trees are deposited on figshare (https://doi.org/10.6084/m9.figshare.26965591.v2, https://doi.org/10.6084/m9.figshare.28381412.v1, https://doi.org/10.6084/m9.figshare.28014992.v1).

## Computer code

There are no previously unreported custom computer codes in this study.

## Acknowledgements

We thank Akinori Yabuki and Kazuo Inaba for providing data and expertise. This work was funded by the Ramón y Cajal Programme (Grant RYC2022-035282-I, funded by the MCIU/AEI/10.13039/501100011033 and the FSE+ to L.J.G.), the Israel Science Foundation (Research Center grants 3131/20 and 1207/24 to O.B.), the European Commission, under Horizon Europe’s research and innovation programme (Bluetools project, Grant Agreement No 101081957 to O.B.), the Nancy and Stephen Grand Technion Energy Program (GTEP), JSPS KAKENHI Grants-in-Aid (grants JP23H04404 to K.I., JP23H04863 to T.N.), JST CREST (grants JPMJCR22N2 to K.I.), and MEXT Promotion of Development of a Joint Usage/ Research System Project: Coalition of Universities for Research Excellence Program (CURE) (grant JPMXP1323015482 to K.I.). O.B. holds the Louis and Lyra Richmond Chair in Life Sciences.

## Ethics declaration

The authors declare no competing interest.

