## Supplementary Information for "Apusomonad rhodopsins, a new family of ultraviolet to blue light absorbing rhodopsin channels"

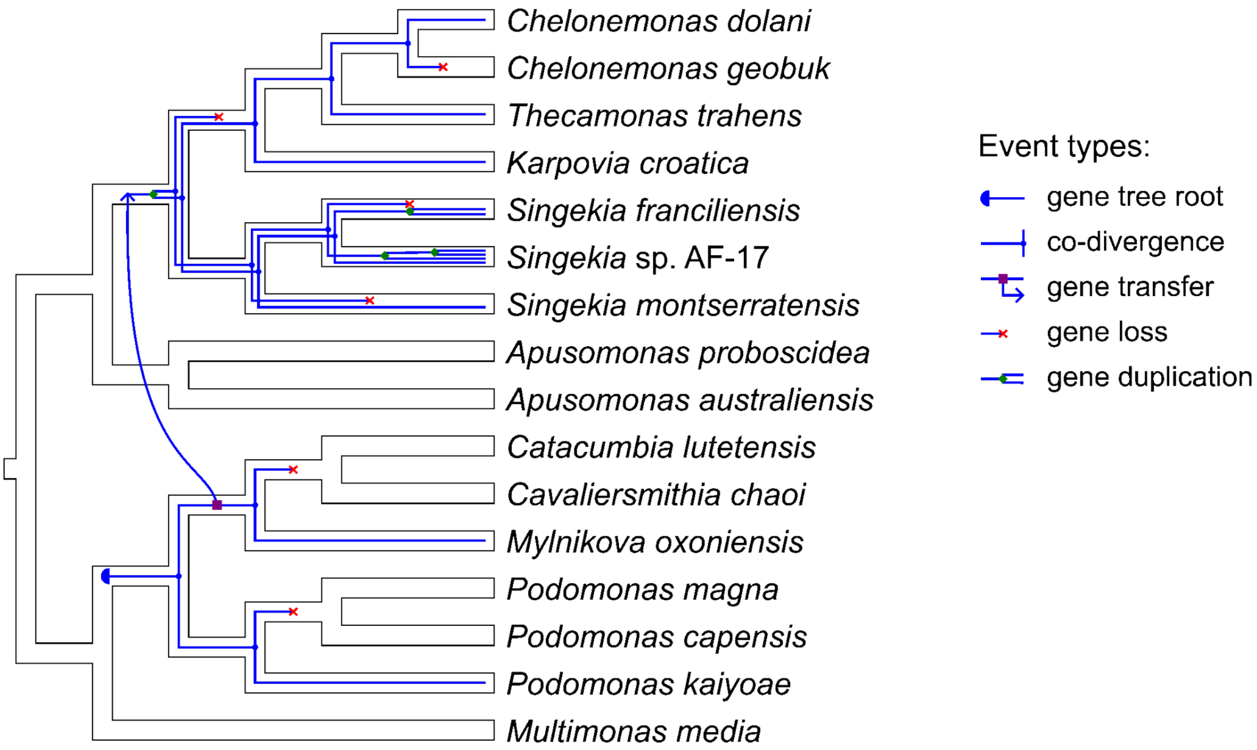


**Figure S1. Evolution of ApuRs among apusomonads.** Results of gene tree-species tree reconciliation based on the phylogenetic inferences for Apusomonadida and ApuRs (see Figure 1). *Podomonas kaiyoae* was placed on the species tree based on the previously published 18S rRNA phylogenetic analysis [1]. Among the equally parsimonious reconstructions, the accelerated-transformation (ACCTRAN) scenario was preferred. Total event score: 15.0, obtained with the default cost values (duplication: 1.5, transfer: 3.0, co-divergence: 0.0, loss: 1.0).

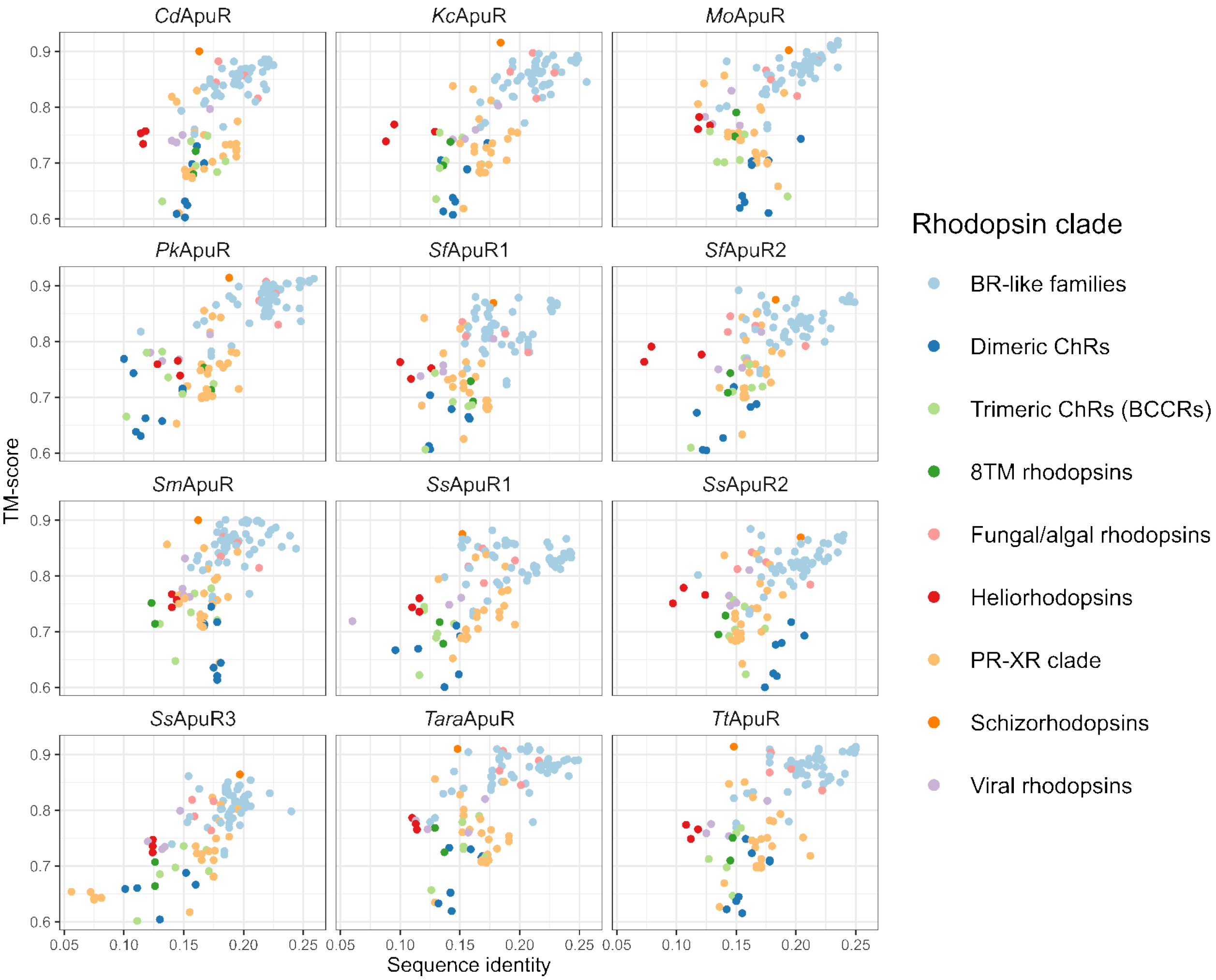


**Figure S2. Similarity between ApuRs and other microbial rhodopsins.** The predicted structures of the ApuR rhodopsin domain were aligned in a pairwise manner with microbial rhodopsin structures available in the Protein Data Bank. Among the multiple structures available for the same reference protein the best-matching one was chosen. The reference structures are colored according to their phylogenetic affiliation.

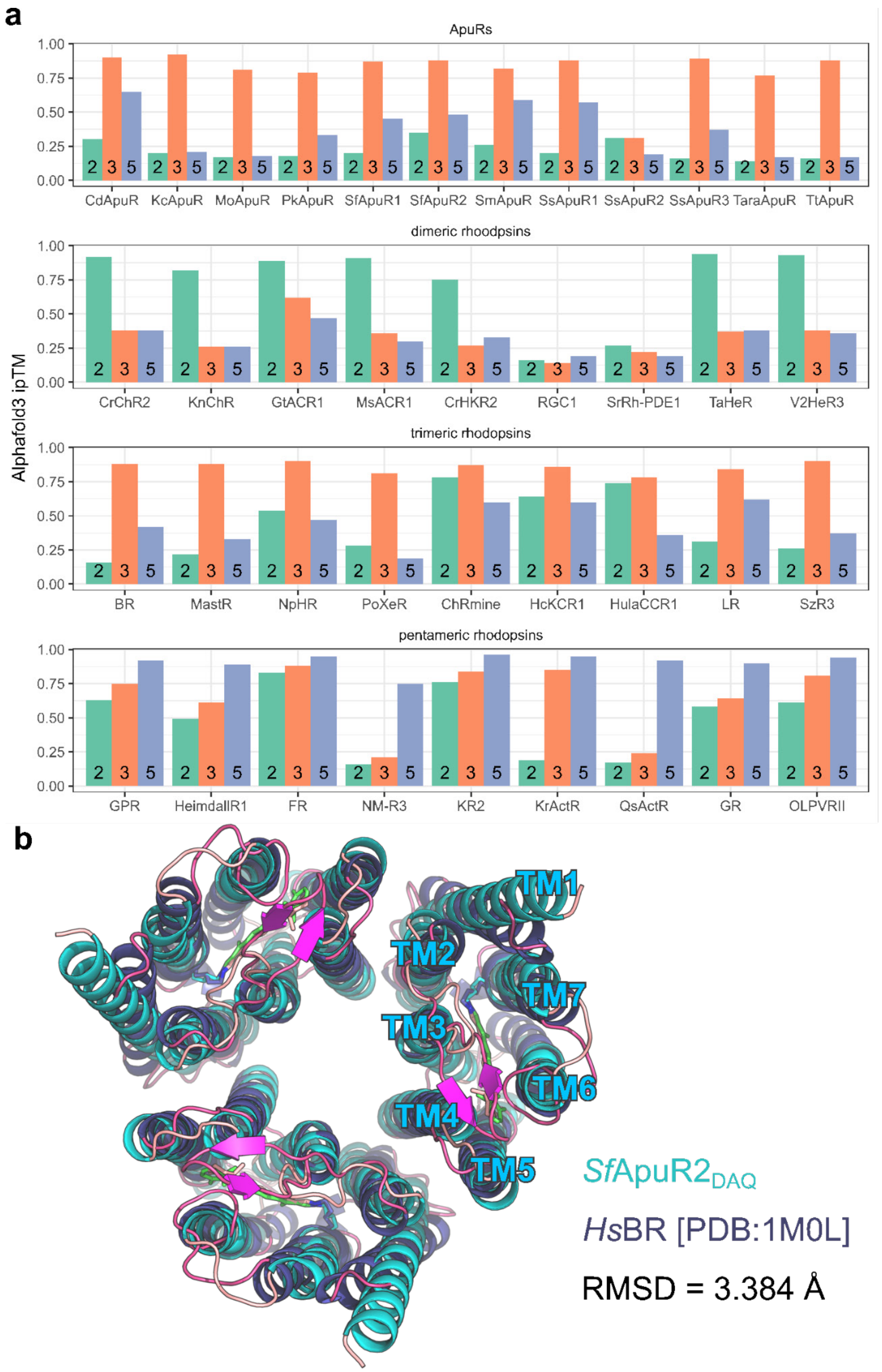


**Figure S3. Prediction of oligomeric structures of ApuRs. a** Estimates of interface prediction accuracy for 2-, 3- and 5-mers of ApuRs and reference rhodopsins with known or predicted oligomeric structures. **b** The predicted trimeric structure of *Sf*ApuR2$_{DAQ}$ in comparison to *Hs*BR, viewed from the extracellular side.

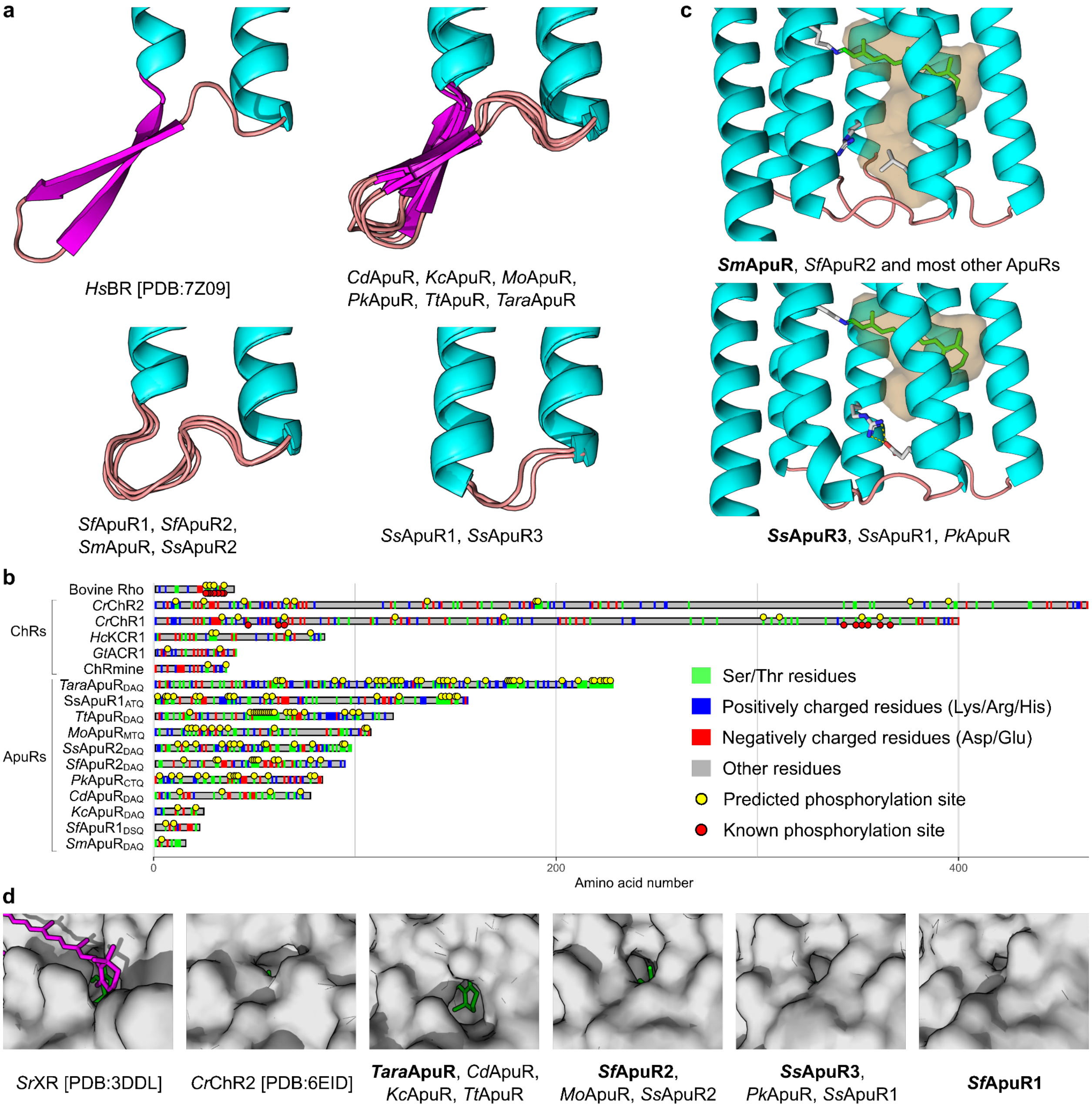


**Figure S4. Additional structural features of ApuRs. a** Variation in the structure of the extracellular loop 1 (ECL1) among ApuRs. ECL1 of *Hs*BR is shown for comparison. **b** C-terminal extensions of ApuRs in comparison to bovine rhodopsin and channelrhodopsins. Indicated are residues of different types, phosphorylation sites predicted with NetPhos, as well phosphorylation sites known for *Cr*ChR1 and bovine rhodopsin. **c** *Sf*ApuR2 position 194 is occupied in some ApuRs by a Glu residue putatively forming a salt bridge with the highly conserved Arg residue R77 (R82 in *Hs*BR) in TM3, which might restrict the connection of the retinal-binding pocket with the extracellular side. **d** Variation in the retinal binding pocket fenestration between helices TM5 and TM6 in ApuRs. Highlighted in bold are representative proteins used to visualize groups of similar fenestrations. Fenestrations in the carotenoid-binding rhodopsin *Sr*XR (the carotenoid antenna salinixanthin shown in magenta) and *Cr*ChR2 are shown for comparison. The retinal moiety can be seen through the fenestrations and is indicated in green.

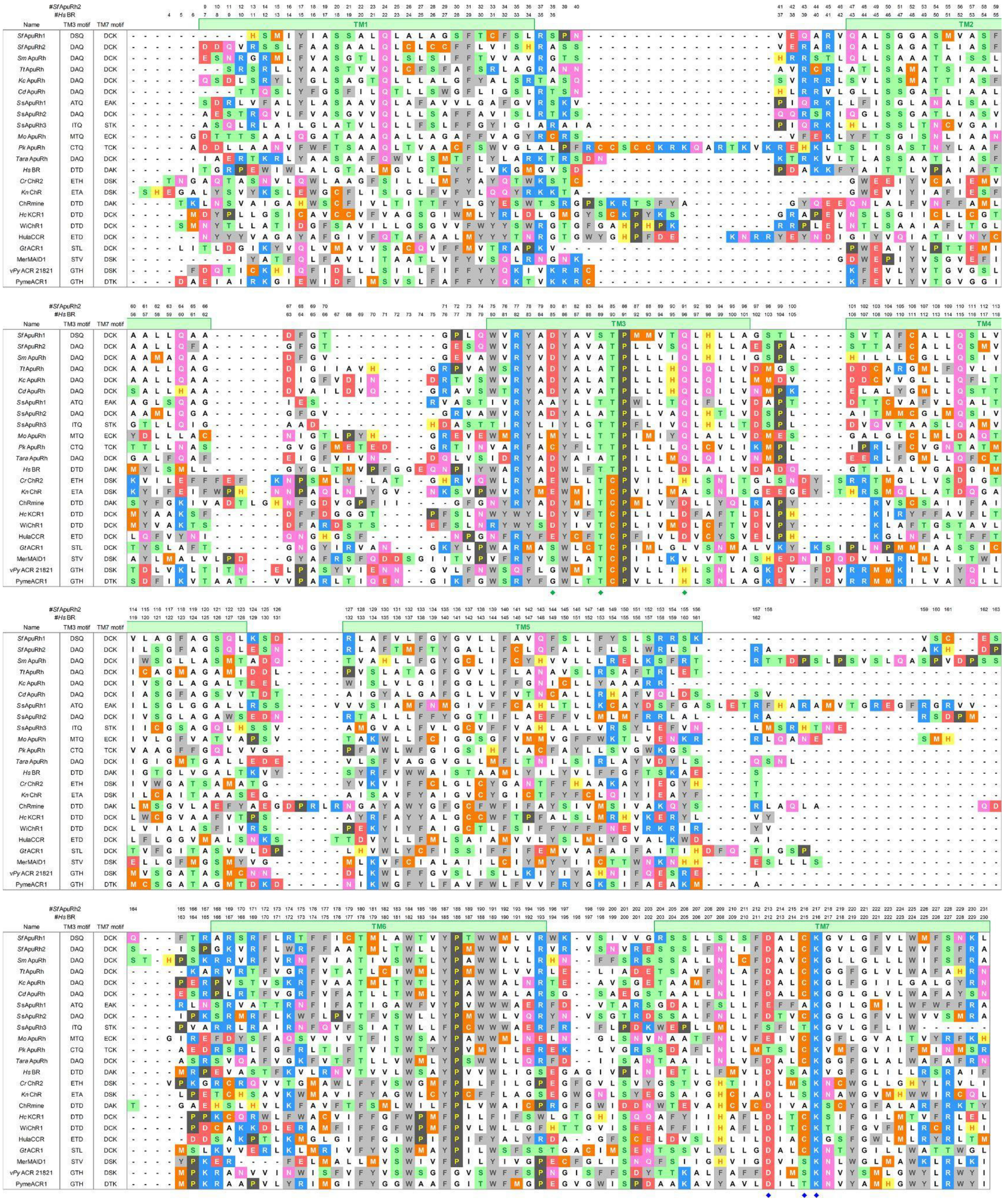

**Figure S5. Amino acid sequence alignments of ApuR with other microbial rhodopsins.** Amino acid sequences of ApuRs are aligned with those of ApuR using PROMALS3D method [2] incorporating three-dimensional structural information of *Hs*BR (PDB ID: 7Z09 [3]), *Cr*ChR2 (PDB

ID: 6EID[4] ), ChRmine (PDB ID: 7W9W [5]), *Hc*KCR1 (PDB ID: 8H86 [6]), and *Gt*ACR1 (PDB ID: 6CSM [7]). The positions of consensus TMs were indicated by green rectangles. The positions of the residues in TM1 and TM2, where the automatic alignment is difficult due to the low conservation of amino acid in the microbial rhodopsin family, were manually corrected based on the three-dimensional structures used in PROMALS3D calculation. The TM3 motif, corresponding to *Hs*BR D85 (counterion), T89, and D96, and the TM7 motif, corresponding to *Hs*BR D212 (counterion), A215, and K216 (binding to the retinal chromophore), are shown in the first and second columns, and their positions are marked with green and blue diamonds, respectively. The amino acids are colored according to their chemical properties: hydrophobic aliphatic residues (white); hydroxyl group-bearing aliphatic residues (green); acidic residues (red); basic residues (blue); Asn and Gln (pink); sulfur-containing residues (orange); proline (black); aromatic residues (gray).

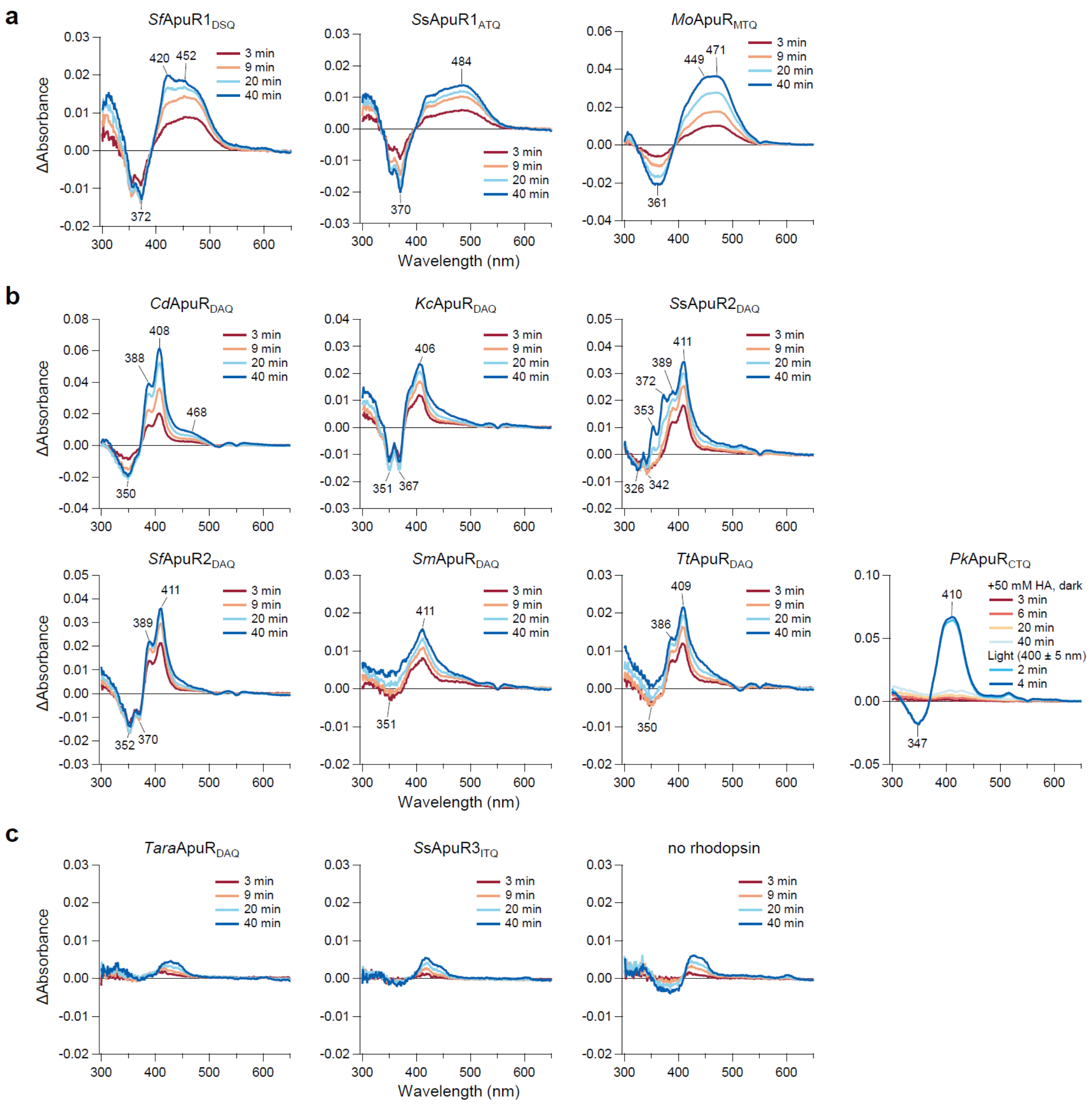


**Figure S6. Hydroxylamine bleaching of ApuRs. a**, **b** Difference spectra of ApuRs that exhibited their main absorption peaks in the blue (**a**) and violet (**b**) regions upon hydroxylamine bleach reaction. Difference spectra for each sample were calculated by subtracting the second-scanned and following spectra from the first-scanned spectrum. To accelerate the bleaching reaction, *Pk*ApuR$_{CTQ}$ was illuminated with light (400 ± 5 nm) after dark measurements. **c** Difference spectra of *Tara*ApuR$_{DAQ}$, *S*sApuR3$_{ITQ}$, and a negative control, no rhodopsin (no plasmid DNA transfected). *Tara*ApuR$_{DAQ}$ and *S*sApuR3$_{ITQ}$ showed only small spectral changes similar to the negative control.

The 420-nm peak of *Sf*ApuR1$_{DSQ}$ is probably due to endogenous cytochrome in COS-1 cells as observed in the negative control (**c**).

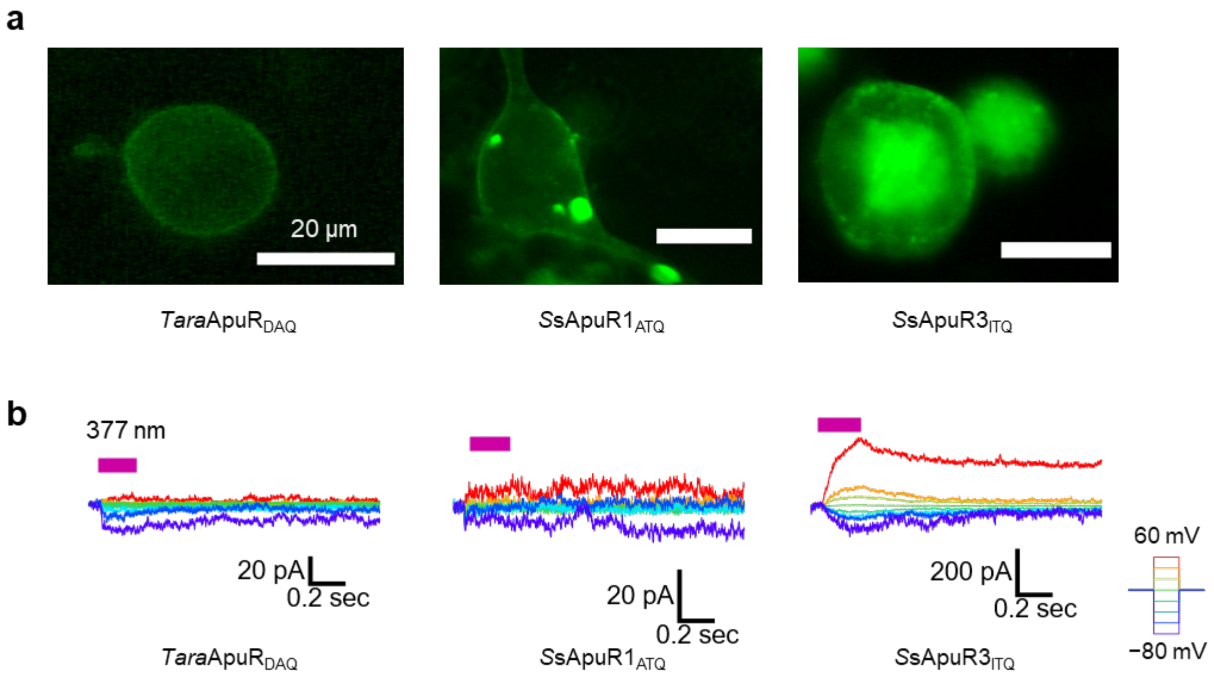


**Figure S7. Some ApuRs exhibit weak photocurrent. a** Fluorescent images of ND7/23 cells expressing *Tara*ApuR$_{DAQ}$, *S*sApuR1$_{ATQ}$, and *S*sApuR3$_{ITQ}$ which are C-terminally labeled with eYFP (Scale bar = 20 µm). **b** Photocurrent traces of *Tara*ApuR$_{DAQ}$, *S*sApuR1$_{ATQ}$, and *S*sApuR3$_{ITQ}$ recorded with standard pipette and extracellular solutions.

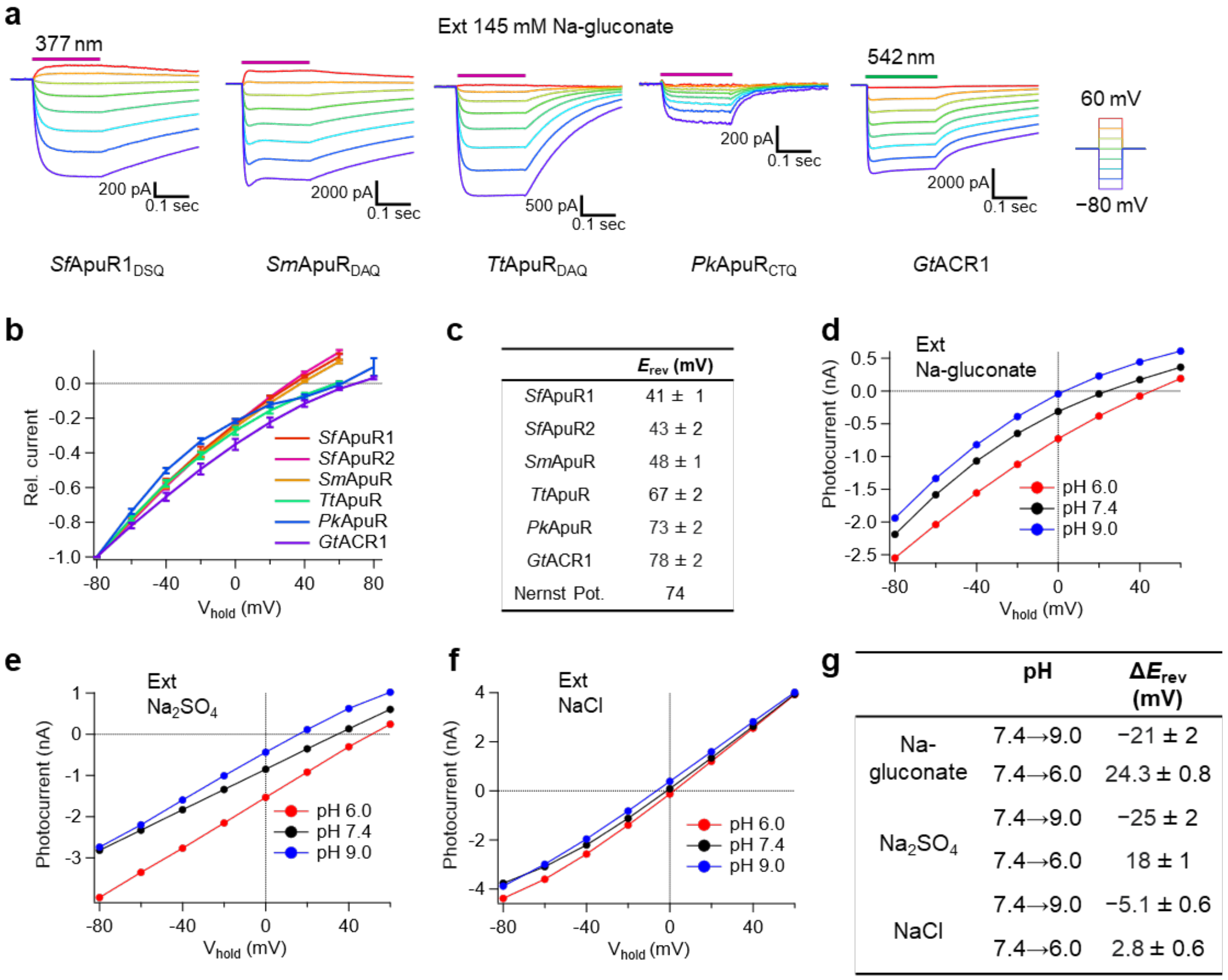


| | $E_{rev}$ (mV) |
|---|---|
| *Sf*ApuR1 | 41 ± 1 |
| *Sf*ApuR2 | 43 ± 2 |
| *Sm*ApuR | 48 ± 1 |
| *Tt*ApuR | 67 ± 2 |
| *Pk*ApuR | 73 ± 2 |
| *Gt*ACR1 | 78 ± 2 |
| Nernst Pot. | 74 |

| | pH | $\Delta E_{rev}$ (mV) |
|---|---|---|
| Na-gluconate | 7.4→9.0 | −21 ± 2 |
| | 7.4→6.0 | 24.3 ± 0.8 |
| $Na_2SO_4$ | 7.4→9.0 | −25 ± 2 |
| | 7.4→6.0 | 18 ± 1 |
| NaCl | 7.4→9.0 | −5.1 ± 0.6 |
| | 7.4→6.0 | 2.8 ± 0.6 |

**Figure S8. Anion permeability of ApuRs and pH dependence of photocurrent of *Sf*ApuR2$_{DAQ}$.** **a** Photocurrent traces of ApuRs and *Gt*ACR1 at 145 mM [Na-gluconate]$_{ext}$ and 130 mM [NaCl]$_{int}$. **b** *I*–*V* curves of ApuRs and *Gt*ACR1 steady current recorded with 145 mM Na-gluconate extracellular and standard pipette solutions (mean ± S.E., $n$ = 4–6, Photocurrents are normalized at −80 mV.). **c** $E_{rev}$ values determined from the *I*–*V* curves shown in **b** and the calculated $Cl^-$ Nernst potential of this experimental condition. $E_{rev}$ values were corrected by the liquid junction potential (mean ± S.E., $n$ = 4–6). **d**–**f** Representative *I*–*V* curves of the *Sf*ApuR2$_{DAQ}$ steady photocurrent (mean current during the latter half of the illumination time) recorded by exchanging extracellular solution with different pH at 145 mM [Na-gluconate]$_{ext}$ (**d**), 95 mM [$Na_2SO_4$] (**e**) and 145 mM [NaCl]$_{ext}$ (**f**). (**g**) $E_{rev}$ shift ($\Delta E_{rev}$) by changing pH of extracellular solutions (mean ± S.E., $n$ = 5).

a

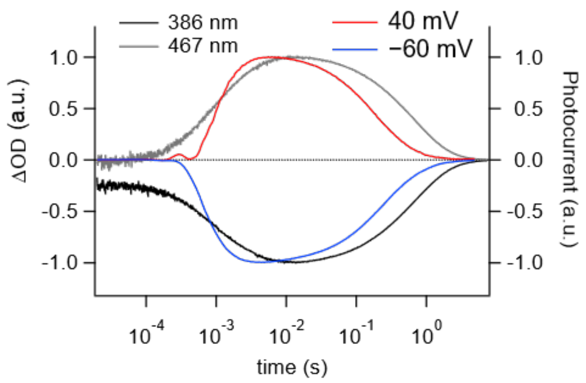


b

| | $Open_1$ / ms (contribution) | $Open_2$ / ms (contribution) |
|---|---|---|
| 40mV | 0.55 ± 0.03 (93.6 $_{\pm 0.4}$%) | 3.4 ± 0.5 (6.4 $_{\pm 0.4}$%) |
| −60mV | 0.48 ± 0.02 (87 $_{\pm 1}$%) | 10 ± 1 (13 $_{\pm 1}$%) |

| | $Close_1$ / ms (contribution) | $Close_2$ / ms (contribution) |
|---|---|---|
| 40mV | 159 ± 7 (65 $_{\pm 2}$%) | 560 ± 50 (35 $_{\pm 2}$%) |
| −60mV | 211 ± 8 (65 $_{\pm 1}$%) | 720 ± 20 (35 $_{\pm 1}$%) |

c

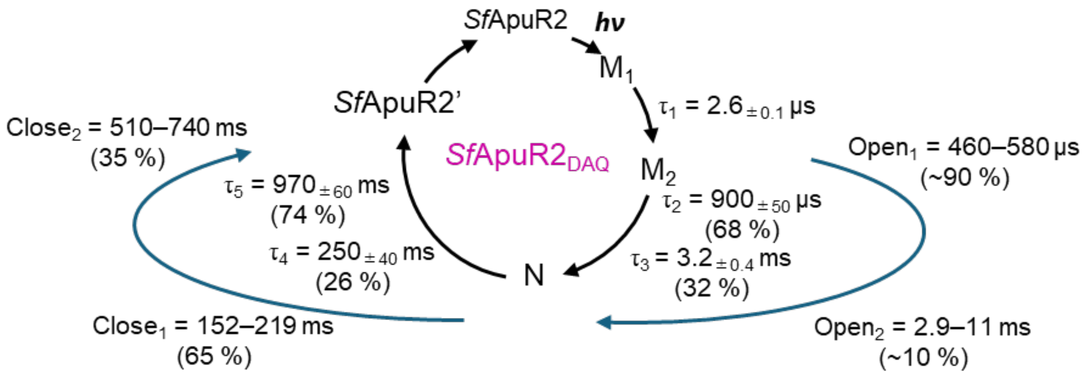


**Figure S9. Comparison of the open/close kinetics with the absorption change of *Sf*ApuR2$_{DAQ}$.** **a** Time traces of absorption changes (386 and 467 nm) of *Sf*ApuR2$_{DAQ}$ and photocurrent traces upon nanosecond laser flash excitation at 40- and −60-mV holding potential (red and blue lines, respectively). Traces were normalized at a peak position. **b** Time constants of channel opening and closing of *Sf*ApuR2$_{DAQ}$ (mean ± S.E., $n$ = 8). **c** The photocycle model of *Sf*ApuR2$_{DAQ}$ including spectroscopically observed intermediates (black arrows) and the channel open/close kinetics (blue arrows).

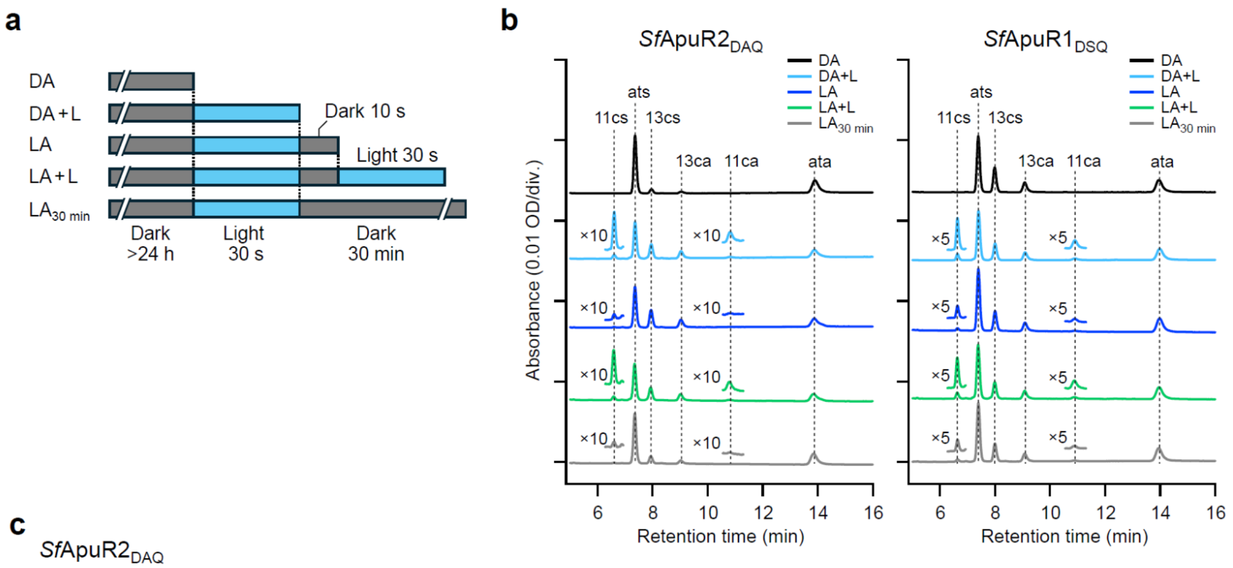


c

*Sf*ApuR2$_{DAQ}$

| | 11-*cis* (%) | 13-*cis* (%) | all-*trans* (%) |
|---|---|---|---|
| DA | 0.27 ± 0.09 | 7.46 ± 0.05 | 92.26 ± 0.07 |
| DA+L | 8.33 ± 0.32 | 29.82 ± 0.54 | 61.85 ± 0.63 |
| LA | 0.86 ± 0.07 | 33.01 ± 0.33 | 66.13 ± 0.39 |
| LA+L | 8.84 ± 0.20 | 27.85 ± 0.07 | 63.31 ± 0.13 |
| LA$_{30\ min}$ | 1.06 ± 0.03 | 14.88 ± 0.99 | 84.06 ± 0.99 |

*Sf*ApuR1$_{DSQ}$

| | 11-*cis* (%) | 13-*cis* (%) | all-*trans* (%) |
|---|---|---|---|
| DA | 0.39 ± 0.01 | 33.96 ± 0.10 | 65.65 ± 0.09 |
| DA+L | 11.31 ± 0.34 | 26.32 ± 0.19 | 62.37 ± 0.18 |
| LA | 3.94 ± 0.16 | 26.59 ± 0.11 | 69.47 ± 0.06 |
| LA+L | 11.14 ± 0.46 | 24.34 ± 0.25 | 64.52 ± 0.22 |
| LA$_{30\ min}$ | 3.39 ± 0.04 | 25.83 ± 0.06 | 70.77 ± 0.02 |

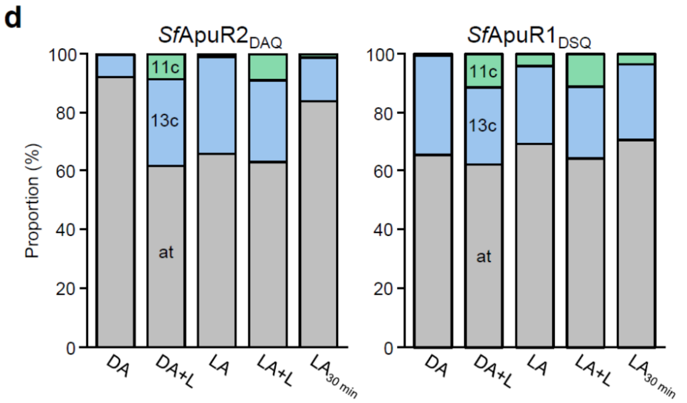


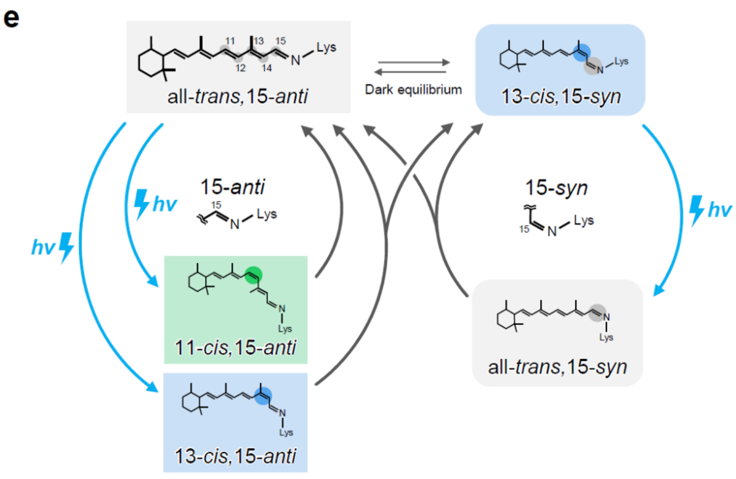


**Figure S10. HPLC analysis of chromophore configurations. a** The light conditions for each sample of HPLC analysis. Purified samples were dark-adapted (DA) for at least 24 h at 4 °C and treated with different light conditions: 30-s illumination (DA+L); DA+L followed by 10-s incubation in the dark (light-adapted, LA); LA followed by 30-s illumination (LA+L); DA+L followed by 30-min incubation in the dark ($LA_{30\ min}$). Illumination (460 ± 5 nm for *Sf*ApuR1$_{DSQ}$; 400 ± 5 nm for *Sf*ApuR2$_{DAQ}$) was performed at room temperature. **b** HPLC patterns of retinal-oxime isomers for *Sf*ApuR2$_{DAQ}$ (left) and *Sf*ApuR1$_{DSQ}$ (right). **c** The composition of the retinal isomer determined by the HPLC analysis ($n$ = 3, mean ± S.E). The molar composition was calculated with the molar extinction coefficients at 360 nm: all-*trans*-15-*syn*: 54,900 $M^{-1}$ $cm^{-1}$; all-*trans*-15-*anti*: 51,600 $M^{-1}$ $cm^{-1}$; 13-*cis*-15-*syn*, 49,000 $M^{-1}$ $cm^{-1}$; 13-*cis*-15-*anti*, 52,100 $M^{-1}$ $cm^{-1}$; 11-*cis*-15-*syn*, 35,000 $M^{-1}$ $cm^{-1}$; 11-*cis*-15-*anti*, 29,600 $M^{-1}$ $cm^{-1}$. **d** The isomer composition in different light conditions. **e** A hypothetical model of the chromophore isomerization in *Sf*ApuRs. 11c: 11-*cis;* 13c: 13-*cis*; at: all-*trans*; 11cs: 11-*cis*, 15-*syn*; 13cs: 13-*cis*, 15-*syn*; ats: all-*trans*, 15-*syn*; 11ca: 11-*cis*, 15-*anti*; 13ca: 13-*cis*, 15-*anti*; ata: all-*trans*, 15-*anti.*

**Table S1. Composition of pipette and extracellular solutions used for ion-selectivity measurements.** All concentrations are in mM. Liquid junction potentials (LJPs) between the NaCl pipette and bath solutions are listed.

| | NaCl pipette | $Cl^-$ ext | $Br^-$ ext | $NO_3^-$ ext | gluconate ext | $SO_4^{2-}$ ext |
|---|---|---|---|---|---|---|
| HEPES | 10 | 10 | 10 | 10 | 10 | 10 |
| NaCl | 130 | 145 | — | — | — | — |
| NaBr | — | — | 145 | — | — | — |
| $NaNO_3$ | — | — | — | 145 | — | — |
| Na gluconate | — | — | — | — | 145 | — |
| $Na_2SO_4$ | — | — | — | — | — | 95 |
| $MgCl_2$ | 1 | 1 | 1 | 1 | 1 | 1 |
| $CaCl_2$ | — | 2.5 | 2.5 | 2.5 | 2.5 | 2.5 |
| $Na_2EGTA$ | 5 | — | — | — | — | — |
| MgATP | 2.5 | — | — | — | — | — |
| glucose | — | 11 | 11 | 11 | 11 | 11 |
| LJP (mV) | — | 2.2 | 2.6 | 1.3 | −9.9 | −4.6 |
